# Genomic prediction of symbiotic interactions between two *Endozoicomonas* clades and their coral host, *Acropora loripes*

**DOI:** 10.1101/2025.02.16.638547

**Authors:** Cecilie R Gotze, Kshitij Tandon, Gayle K Philip, Ashley M Dungan, Justin Maire, Lone Hoj, Linda L Blackall, Madeleine JH van Oppen

**Affiliations:** School of BioSciences, The University of Melbourne, Parkville, VIC 3010, Australia; Australian Institute of Marine Science, Townsville, QLD 4810, Australia; Melbourne Integrative Genomics, The University of Melbourne, Parkville, VIC 3010, Australia; Department of Microbiology and Immunology at the Peter Doherty Institute of Infection and Immunity, The University of Melbourne, Parkville, VIC 3010, Australia; Melbourne Bioinformatics, The University of Melbourne, Parkville, VIC 3010, Australia

**Keywords:** environmental microbiology, coral, symbiosis, microbiome, *Endozoicomonas*, comparative genomics, phylogenomics

## Abstract

**Background:** The bacterial genus *Endozoicomonas* is a predominant member of the coral microbiome, widely recognised for its ubiquity and ability to form high-density aggregates within coral tissues. Hence, investigating its metabolic interplay with coral hosts offers critical insights into its ecological roles and contributions to coral health and resilience.

**Results:** Using long- and short-read whole-genome sequencing of 11 *Endozoicomonas* strains from *Acropora loripes*, genome sizes were found to range between 5.8 and 7.1 Mbp. Phylogenomic analysis identified two distinct clades within the family Endozoicomonadaceae. Metabolic reconstruction uncovered clade-specific pathways, including the degradation of holobiont-derived carbon and lipids (e.g., galactose, starch, triacylglycerol, D-glucuronate), latter of suggest involvement of *Endozoicomonas* in host ‘sex-type’ steroid hormone metabolism. A clade-specific type 6 Secretion System (T6SS) and predicted effector molecules were identified, potentially facilitating coral-bacterium symbiosis. Additionally, genomic analyses revealed diverse phosphorus acquisition strategies, implicating *Endozoicomonas* in holobiont phosphorus cycling and stress responses.

**Conclusions:** This study reveals clade-specific genomic signatures of *Endozoicomonas* supporting its mutualistic lifestyle within corals. Findings suggests possible roles in nutrient cycling, reproductive health, and stress resilience, offering novel insights into coral holobiont functioning and potential strategies for reef restoration.

## Background

Scleractinian corals support a highly diverse community of symbiotic microorganisms, including dinoflagellate photosymbionts (Symbiodiniaceae), bacteria, archaea, viruses, and fungi, collectively known as the coral holobiont ^[1–4]^. While the coral-Symbiodiniaceae mutualism has been extensively studied ^[5,6]^, less is known about coral-bacteria interactions, which can range from mutualistic to antagonistic ^[3,7]^. Bacterial communities exhibit significant variability in diversity across coral species, influenced by degrees of host specificity and environmental disturbances ^[8,9]^. In recent years, these bacterial communities have garnered increased attention for their potential contributions to holobiont functioning and health, including nutrient cycling, immune modulation, and pathogen resistance ^[10–12]^. Among the coral-associated bacteria, the genus *Endozoicomonas* stands out due to its association with a wide range of marine hosts, including sponges, sea anemones, ascidians and some fish ^[13–18]^. Metagenomic and transcriptomic data suggest that *Endozoicomonas* is metabolically versatile, which may explain its broad spectrum of interactions with a diverse range of marine invertebrates, spanning from mutualistic to commensal relationships ^[19]^. Furthermore, *Endozoicomonas* are frequently detected in high relative abundance in coral microbiomes, suggesting a potentially important ecological role in coral health and functioning. However, while they are consistently found across a diverse range of coral taxa and geographic regions, they are not necessarily specific to any one species ^[7,20–22]^.

Within coral tissues, *Endozoicomonas* commonly forms densely populated clusters known as cell-associated microbial aggregates (CAMAs), suggesting adaptation to specific microenvironments within the coral colony ^[23–25]^. Their known functional roles include sulfur cycling and the potential transfer of amino acids synthesised *de novo*, presumably for the benefit of its hosts ^[26,27]^. A study on *Stylophora pistillata* revealed that CAMAs are enriched with phosphorus, with *Endozoicomonas* cells showing elevated phosphorus levels compared to coral tissues and photosymbiotic Symbiodiniaceae. This suggests that *Endozoicomonas* may play a significant role in sequestering and cycling phosphate within the coral holobiont ^[24]^. Considering the oligotrophic nature of coral reef waters, the amount of phosphorous found within high-density CAMAs likely contributes significantly to the overall phosphorous biogeochemistry of the holobiont ^[28]^. Given the importance of phosphorus in both microbial and coral metabolism, this observation implies that *Endozoicomonas* may play a key role in regulating nutrient availability within the holobiont ^[29–31]^. Despite these insights, our understanding of how the diversity of *Endozoicomonas* species contributes to coral biochemical pathways is limited. Hence, more research is needed to elucidate the mechanistic interactions between *Endozoicomonas* and its hosts, particularly the specific functional roles different strains or species may play in coral health and resilience.

A previous study recently described the phylogenetic placement of 11 *Endozoicomonas* strains isolated from the common reef-building coral *Acropora loripes* in two distinct clades (Clade-A and Clade-B) based on 16S rRNA gene sequences ^[32]^. The spatial distribution of clade members within the host was also mapped using clade-specific fluorescence *in situ* hybridization probes and confocal microscopy, with CAMAs consistently observed within the gastrodermal cell layer, near the host photosymbionts. These CAMAs exhibited clade-specific morphologies where members of Clade-A formed structured aggregates with a clear boundary, whereas members of Clade-B displayed unrestricted growth and formed clusters lacking a clear boundary. This disparity prompted the investigation into whether the differing aggregation patterns correspond to differences in genomic features of Clade-A and -B strains and in the type of symbiosis they can form with their coral host. Therefore, in this study, we sequenced the 11 *Endozoicomonas* strains isolated in ^[32]^ and carried out comparative genomic analyses to elucidate 1) the mechanisms by which *Endozoicomonas* potentially acquire and transfer nutrients to and from other holobiont members (*i.e.,* coral or Symbiodiniaceae), 2) underlying mechanisms potentially governing aggregation patterns, and 3) mechanisms by which *Endozoicomonas* can cope with environmental challenges, such as nutrient deprivation and the host immune response. These data are discussed in the context of the putative physiological role each of the clades plays within the coral holobiont.

## Results and Discussion

### Genomic assembly profiles

Whole-genome sequencing of 11 *Endozoicomonas* isolates obtained from the coral *A. loripes* showed that all assemblies had high completeness of 99.1 ± 0.1% and low levels of contamination (1.3 ± 0.7%) with contig numbers ranging from 1-413. Guanine-cytosine (GC) content averaged at 49.3 ± 0.1% for Clade-A and 47.8 ± 0.1% for Clade-B. All genomes were relatively large (Table 1), with high coding densities compared to the smallest genome within Endozoicomonadaceae. Clade-A generally had smaller genomes whereas Clade-B had larger genomes with slightly higher coding densities (Clade-A: 6.01 ± 0.2 Mbp, 84.95 ± 0.9% and Clade-B: 7.01 ± 0.1 Mbp and 85.7 ± 1.5%; Table 1, Fig. 1). This corroborates the notion that *Endozoicomonas* spp. are not obligate symbionts, which typically have smaller and more streamlined genomes compared to bacteria with a free-living stage ^[34]^.

**Fig. 1.**
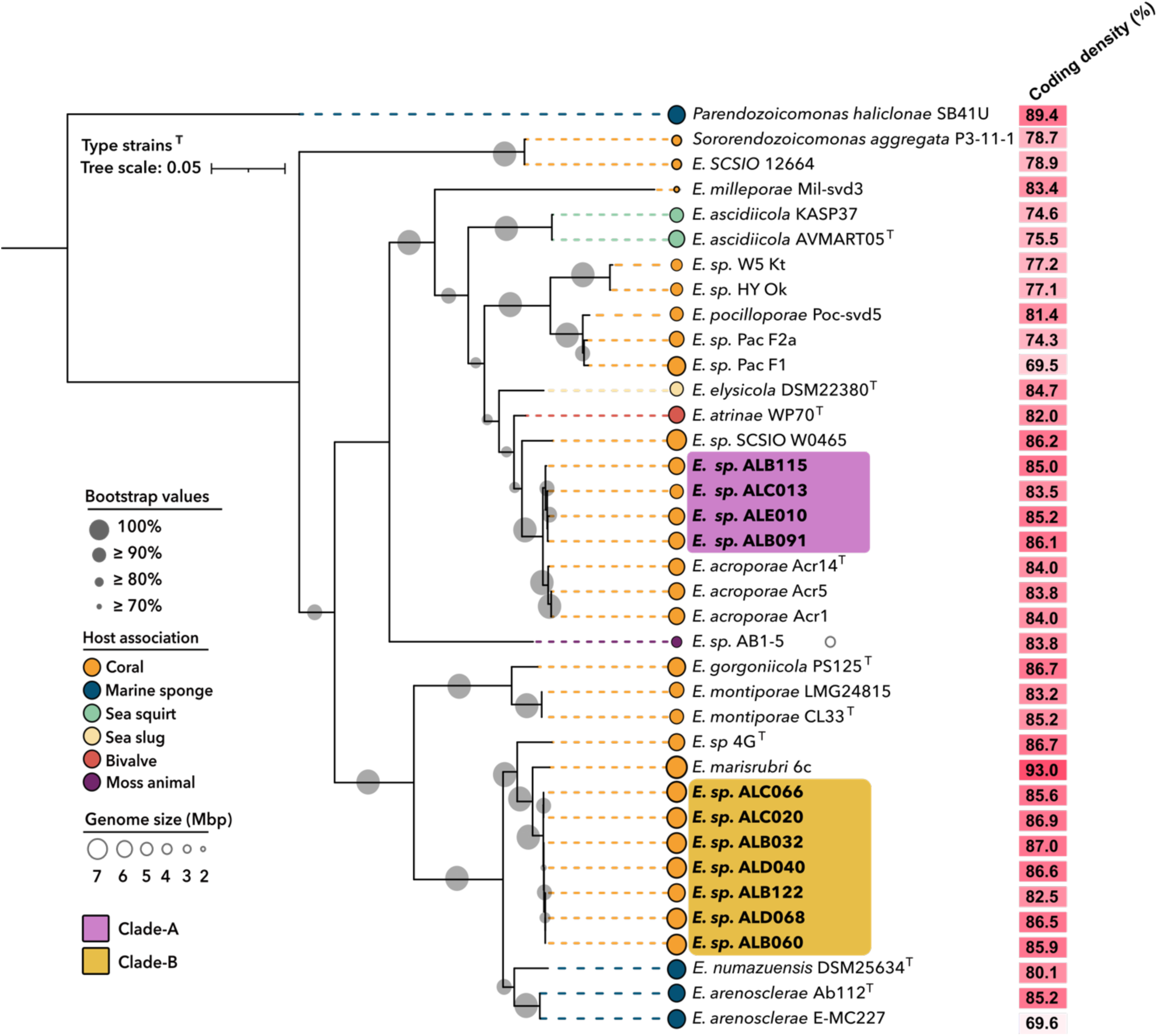
Whole genome phylogenetic reconstruction of Endozoicomonadaceae. Maximum likelihood phylogenetic tree based on 120 bacterial marker genes. Isolates sequenced in this study are shown in boldface. The tree is rooted at *Parendozoicomonas haliclonae* S-B4-, with *Sororendozoicomonas aggregata* Pac-P3-11-1 and *Endozoicomonas* SCSIO 12664 as outgroups. Bootstrap support values are provided based on 1000 replicates. *Endozoicomonas* strains isolated from different invertebrate hosts are colour-coded according to their host isolation source.

**Table 1.** Assembly statistics obtained from 11 pure cultures of *Endozoicomonas* isolated from *Acropora loripes*.

| Genome | Genus assigned by GTDB-Tk | Genome Size (bp) | Completeness (%) | Contamination (%) | G+C content (%) | N50 | L50 | Contigs | Number of: |  |  |
| --- | --- | --- | --- | --- | --- | --- | --- | --- | --- | --- | --- |
|  |  |  |  |  |  |  |  |  | CDSs | rRNAs | tRNAs |
| Clade A |  |  |  |  |  |  |  |  |  |  |  |
| ALE010 | *Endozoicomonas* | 6,014,084 | 98.99 | 2.26 | 49.30 | 73,017 | 25 | 198 | 4925 | 8 | 75 |
| ALC013 | *Endozoicomonas* | 5,797,801 | 98.83 | 2.24 | 49.19 | 30,792 | 57 | 413 | 4779 | 1 | 72 |
| ALB091 | *Endozoicomonas* | 6,211,304 | 98.99 | 2.20 | 49.37 | 1,189,902 | 3 | 21 | 5001 | 23 | 84 |
| ALB115 | *Endozoicomonas* | 6,016,619 | 98.99 | 1.77 | 49.29 | 75,305 | 23 | 206 | 4977 | 7 | 79 |
| Clade B |  |  |  |  |  |  |  |  |  |  |  |
| ALC020 | *Endozoicomonas* | 7,035,133 | 99.07 | 0.54 | 47.82 | 7,035,133 | 1 | 1 | 5008 | 21 | 83 |
| ALB032 | *Endozoicomonas* | 7,138,852 | 99.02 | 0.97 | 47.83 | 7129341 | 1 | 2 | 5105 | 21 | 82 |
| ALD040 | *Endozoicomonas* | 7,102,962 | 99.07 | 1.83 | 47.85 | 7082469 | 1 | 2 | 5034 | 21 | 82 |
| ALB060 | *Endozoicomonas* | 7,038,320 | 99.07 | 0.54 | 47.81 | 2,395,980 | 2 | 143 | 4980 | 21 | 81 |
| ALC066 | *Endozoicomonas* | 7,007,201 | 99.07 | 0.75 | 47.75 | 677,632 | 4 | 64 | 5358 | 19 | 81 |
| ALD068 | *Endozoicomonas* | 7,013,663 | 99.07 | 0.54 | 47.83 | 6,982,335 | 1 | 5 | 4969 | 21 | 82 |
| ALB122 | *Endozoicomonas* | 6,695,625 | 99.07 | 0.75 | 47.71 | 35,739 | 58 | 389 | 4935 | 4 | 79 |

A maximum-likelihood phylogeny based on 120 bacterial marker genes corroborated the 16S rRNA gene phylogeny previously reported ^[32]^. Genome assemblies clustered into two distinct phylogroups, previously designated as Clade-A and Clade-B (Fig. 1). Four isolates clustered together in Clade-A, to which the closest relatives are members of *E. acroporae* isolated from an undocumented *Acropora* sp. ^[35]^. These four strains are also closely related to *Endozoicomonas sp.* SCSIO W0465 isolated from the gastric cavity of the stony coral *Galaxea fascicularis* ^[36]^. The remaining seven isolates formed Clade-B, which clustered with *E. marisrubi* 6c isolated from *Acropora humilis* ^[27]^ and *Endozoicomonas* sp. 4G (obtained from *Acropora muricata* ^[17]^. Members of Clade-A had an average of 95.3% ANI and 60.2 GGD with their closest neighbour *E. acroporae* Acr-12, whereas members of Clade-B exhibited 91% ANI and 26.6 GGD with their closest neighbour *E. marisrubi 6c* (Fig S1) and (Fig S2). Based on the phylogenetic tree and the threshold for denoting a new bacterial species within the same genus ^[37]^ which requires a GGD no less than 70% to its closest described representative, both Clade-A and B likely represent previously undescribed species.

Although strains from the two clades were isolated from the same *A. loripes* coral colonies and their general co-occurrence within individual colonies was confirmed by 16S rRNA gene metabarcoding ^[32]^, they exhibited an ANI of only ∼70% (Fig. S1). This trend was further supported by a low AII of ∼23% (Fig. S2) and *in silico* GGD values of ∼68% (Fig. S3), indicating that these groups diverged in the evolutionary distant past. The pattern in which isolated *Endozoicomonas* strains are not grouped by host identity and where divergent species coexist within the same host has been observed in other acroporid and pocilloporid corals ^[16,38]^. This suggests that a diverse set of *Endozoicomonas* species can colonise not only a single coral species but even a single coral colony ^[17,32]^.

### Acquisition and transfer of carbohydrates

Understanding the metabolic relationships between members of the coral holobiont is paramount in deciphering the symbiotic role of aggregate-forming bacteria within corals. Notably, some *Endozoicomonas* phylotypes establish CAMAs within the epidermis of coral tentacles ^[39–41]^, whereas strains used in this study (Clade-A and Clade-B) were consistently found to colonise the gastrodermis of the mesenteries and actinopharynx near Symbiodiniaceae ^[32]^. Consequently, a comprehensive analysis of their genomes was conducted to elucidate metabolic pathways that may underpin differences in formation.

Carbohydrates exist within coral tissues as di-, oligo- or polysaccharides of hexose sugar molecules (*e.g.,* glucose, fructose or galactose) including cellulose and starch. The major monosaccharides, galactose and glucose, are abundant in coral mucus as well as in Symbiodiniaceae photosynthates ^[42–44]^. While glucose readily enters glycolysis, galactose must first be converted into a derivative of glucose. Members in Clade-A were found to have the genetic potential to take up and assimilate galactose, via the Leloir pathway, which allows the catabolism of galactose via conversion to uridine diphosphate galactose (Fig. 2). This is corroborated by the finding that members of Clade-A encodes homologs to a high-affinity galactose transport system, Mgl. Once inside the cell, galactose can either be converted to glucose-1-phosphate via GalU (present in both clades) or enter the Leloir pathway (in Clade-A only), where it is converted to energy either via the glycolytic or Entner-Doudoroff pathway ^[45]^. Galactose utilisation via the Leloir pathway appears to be highly conserved in most reference *Endozoicomonas* genomes and may be a contributing factor for the co-localisation of *Endozoicomonas* CAMAs and Symbiodiniaceae within the coral tissue. It is plausible that strains with high-affinity galactose transport systems would have a competitive advantage for the uptake of galactose against a low concentration gradient. In contrast, members of Clade-B lack three essential genes in the Leloir pathway, suggesting a reliance on alternative carbon sources, e.g. fructose, mannose and xylose ^[46]^. This is further corroborated by the finding that no galactose transport systems were identified in Clade-B. Noteworthy, some bacteria, such as *Escherichia coli*, that are devoid of systems for the active transport of galactose can successfully grow on galactose - albeit only at high concentrations ^[47]^. Thus, Clade-B may be able to import galactose facilitated by diffusion across the cell membrane, however, the genes responsible for the conversion into substrates that can enter glycolysis are lacking ^[35]^.

**Fig. 2.**
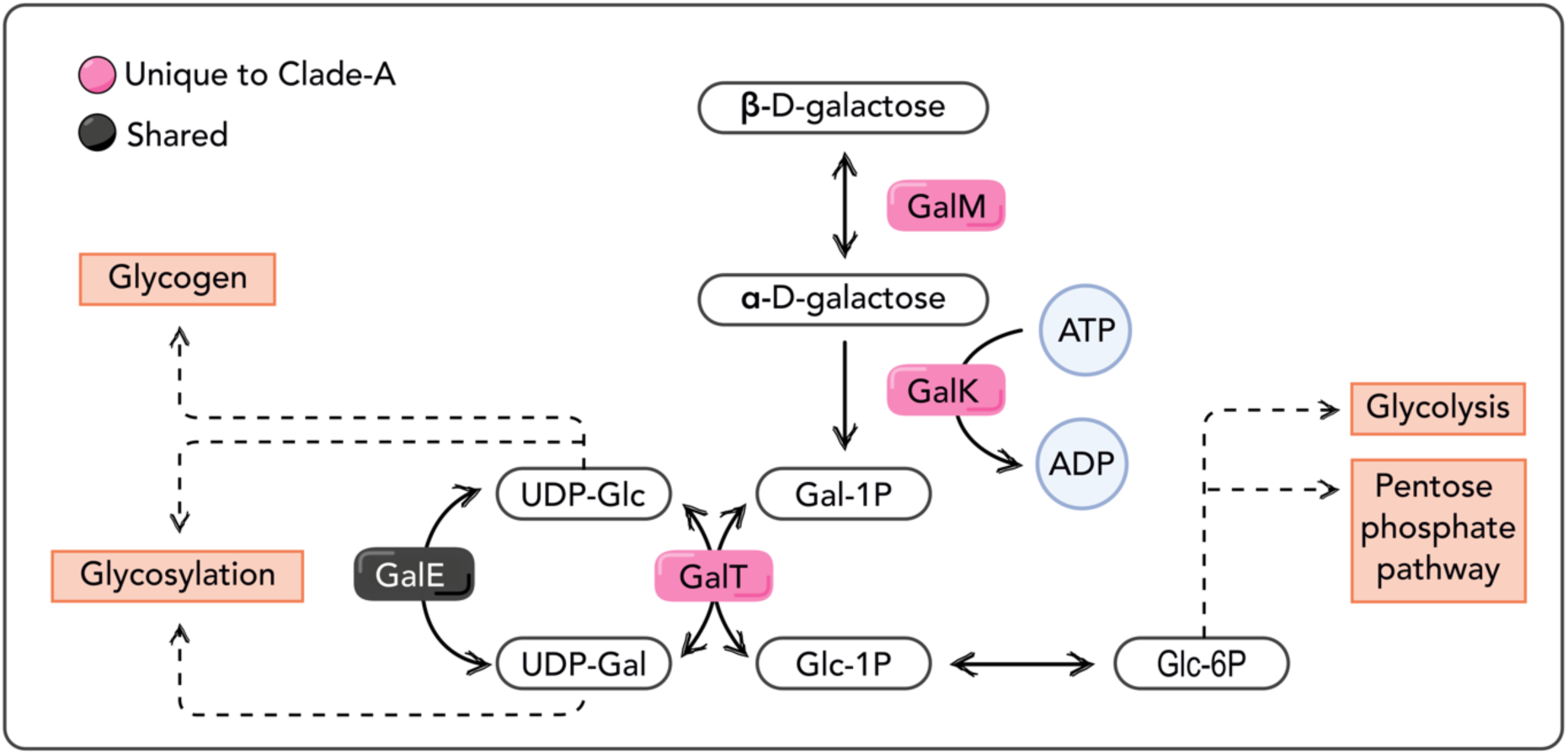
Metabolic diagram illustrating the Leloir pathway in *A. loripes-*associated *Endozoicomonas* for galactose utilisation. β-D-galactose is initially converted to α-D-galactose through the activity of galactose mutarotase (GalM). Subsequently, galactokinase (GalK) catalyzes the phosphorylation of α-D-galactose, utilizing ATP, to produce galactose-1-phosphate (Gal-1P). Gal-1P is then transformed by galactose-1-phosphate uridylyltransferase (GalT), yielding glucose-1-phosphate (Glc-1P) and UDP-galactose (UDP-Gal). The resulting UDP-Gal can be converted to UDP-glucose (UDP-Glc) via the action of UDP-galactose 4’-epimerase (GALE). Glc-1P, derived either directly or through UDP-Glc, enters central metabolic pathways, such as glycolysis through glucose-6-phosphate (Glc-6P) or the pentose phosphate pathway. Dashed arrows in the diagram highlight potential alternative fates of intermediates, including roles in glycogen synthesis or glycosylation.

The absence of genes related to galactose metabolism in Clade-B led us to explore alternative pathways for carbon utilisation. We found that only members of Clade-B possess the complete pathway for taking up and metabolising fructose, a type of dissolved free sugar likely originating from the decomposition of glucans and phytoplankton ingested by the host (Fig. 4). In Clade-B, fructose may be transported into the cell via the fucose transporter FucP, where it can be converted into energy ^[48]^. In contrast, no fructokinases or systems for transporting fructose were found in Clade-A members.

Importantly, Clade-A encodes multiple predicted exoenzymes, including α-Amylase required for the hydrolysis of starch as well as other enzymes degrading complex polysaccharides (*i.e.,* pullulanase), that are missing in Clade-B ^[49,50]^. This suggests that Clade-A has the potential to access stored energy reserves within coral tissues, converting products of photosynthesis such as starch into simple, bioavailable sugars. While both clades possess genes for the assimilation of maltose/maltodextrin, which is the product of starch hydrolysis, via a maltose ABC transporter (Fig. 4), Clade-B would be reliant on exogenous starch/glycogen degradation processes to utilise free maltose.

Taken together, our findings indicate that *Endozoicomonas* species in the two clades have developed distinct strategies to access a shared holobiont resource, namely carbohydrates stored in the coral tissues derived from photosymbionts and/or ingested matter from heterotrophic feeding. Their capacity to utilise various holobiont-derived compounds as carbon sources underscores the metabolic adaptability of this genus to the host environments ^[22]^. This flexibility may confer competitive advantages to *Endozoicomonas* populations within the host environment, enhancing their survival and proliferation ^[19]^. Notably, members of Clade-A and Clade-B occasionally occur as mixed communities within the same CAMAs ^[32]^, indicating the potential for metabolic complementation within these aggregates ^[51]^. This co-occurrence suggests a possible synergistic interaction where Clade-A might break down complex carbohydrates like starch extracellularly, providing simpler sugars such as maltose for both clades. In turn, Clade-B may contribute to the overall metabolic diversity by utilising other carbon sources like fructose. The observation of mixed *Endozoicomonas* phylotypes within the same CAMAs of *Stylophora pistillata* ^[24]^ further supports the hypothesis that co-localisation may facilitate metabolic complementation, enhancing the efficiency of resource cycling within the coral holobiont. Such interactions could promote a more adaptable and resilient symbiotic community capable of thriving in varying environmental conditions ^[52–54]^.

### Degradation of glycosaminoglycans (D-glucuronate degradation)

Clade-A also possess genes for the complete pathway for D-glucuronate degradation, previously undescribed in *Endozoicomonas.* This involves the process by which bacteria break down D-glucuronate, a sugar acid derived from glucose metabolism ^[55]^. This degradation pathway involves several enzymatic steps unique to members of Clade-A, which ultimately convert D-glucuronate (GlcA) into intermediate metabolites that can be further utilised for energy production or biosynthesis (Fig. S4) ^[45]^. The major enzyme of this pathway, β-glucuronidase (GUS), is responsible for cleaving GlcA from either small molecules or the terminal ends of polysaccharides such as glycosaminoglycans (also termed mucopolysaccharides), which are commonly found in coral mucus ^[56]^.

The ability to utilise GlcA would provide the bacteria with the metabolic flexibility to thrive in environments where GlcA is abundant as a carbon source. However, given the predominantly tissue-associated nature of *Endozoicomonas*, this pathway may also be adapted for the utilisation of other holobiont-derived compounds containing GlcA, rather than solely relying on those present in surface mucus ^[24,25,39]^. Notably, GlcA is a common metabolic intermediate in a process known as glucuronidation, which is the major route for the elimination/detoxification of various endogenous and xenobiotic compounds (e.g. heavy metals, oil hydrocarbons, pesticides and herbicides) in a range of vertebrates and invertebrates (Fig. 3) ^[57–59]^. In glucuronidation, GlcA is conjugated to a substrate molecule, typically a lipophilic compound, by enzymes known as UDP-glucuronosyltransferases (UGTs)^[60]^. This conjugation reaction forms a glucuronide conjugate, which is more hydrophilic, effectively increasing the solubility and thereby easier clearance from tissues than the original substrate molecule ^[58,59,61]^.

**Figure 3.**
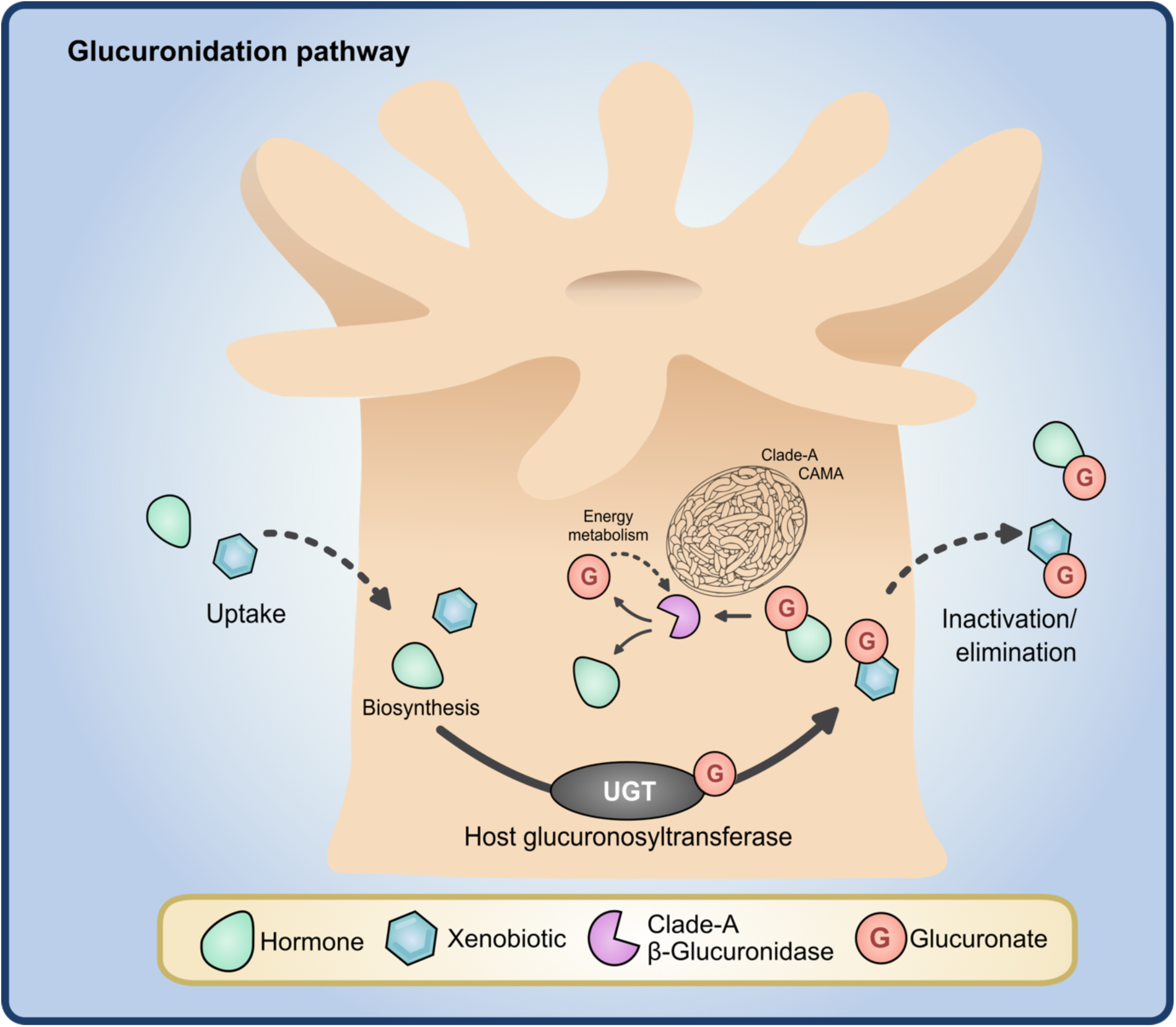
Proposed involvement of *Endozoicomonas* in the host Glucuronidation pathway. Schematic representation of the glucuronidation pathway illustrating the biotransformation of anthropogenic pollutants (xenobiotics) and endogenous compounds (hormones) by host UDP-glucuronosyltransferase (UGT) enzymes. The process involves the transfer of a glucuronic acid moiety from UDP-glucuronate to a substrate molecule, forming a glucuronide conjugate. This conjugation reaction facilitates the detoxification and elimination of hydrophobic compounds from host tissues via the formation of water-soluble glucuronide metabolites. However, coral-associated bacteria with beta-glucuronidase activity have the potential to hydrolyse glucuronide conjugates, leading to the release of the glucuronic acid sugar derivative which they can catabolise for energy production. This process will also liberate the parent compounds influencing the overall clearance of endogenous and exogenous compounds.

While glucuronidation is not well described in cnidarians, major ‘sex-type’ steroid hormones in corals are often found conjugated with GlcA, such as estradiol glucuronide and testosterone glucuronide ^[62]^. Both free and conjugated hormones are consistently detected in coral tissues, with concentrations peaking in both coral tissues and surrounding seawater just before major mass spawning events ^[63,64]^. These hormones are therefore thought to serve as chemical cues for gamete release, as is well-established in vertebrates ^[64]^. Therefore, glucuronidated hormones may represent a significant energy source to bacteria capable of cleavage of the GlcA moiety. As GlcA is highly polar, its conjugation with less polar compounds such as steroid hormones results in more hydrophilic metabolites for easy clearance, thereby decreasing overall hormone levels ^[59]^. Conversely, microbial GlcA removal would extend their retention within coral tissues by returning them to more lipophilic forms ^[65,66]^. While the prevalence and impact of microbial cleavage of glucuronide-conjugated hormones on the coral host are unknown, growing evidence in humans highlights the impact of microbial GUS on host hormone levels. Bacteria capable of GlcA degradation have been implicated in influencing hormone balance in the human body, both directly and indirectly. Directly, they may intercept and cleave GlcA from hormone conjugates, impacting the overall clearance rate of hormones ^[67,68]^. Indirectly, the degradation of GlcA could potentially influence the overall availability of GlcA in the body, a precursor molecule for glucuronidation reactions, which in turn could influence hormone levels and metabolism ^[69–71]^. Recent research has shown the involvement of *Endozoicomonas* species in coral steroid metabolism. For instance, *E. acroporae* (a species closely related to Clade-A, Fig. 1), was able to use the precursor cholesterol both for steroid hormone synthesis (e.g., synthesising progesterone and testosterone) and as a carbon source ^[72]^. Subsequently, this activity was correlated to increased host performance during heat stress ^[72]^. Moreover, another *Endozoicomonas* species, *E. montiporae*, has been shown to metabolise a range of steroids in culture, including estrogen and testosterone. Notably, one of the genes found to be responsible for the conversion of cholesterol *choD*, is located downstream of GUS within Clade-A genomes. This arrangement may have implications for gene regulation, as sequences in the upstream region of one gene can affect the expression of the downstream gene ^[73]^.Additionally, it may suggest that the two genes are part of the same genetic pathway or operon, where they work together to perform a specific function ^[74–76]^.

As steroid glucuronides are biologically less reactive than their parent steroids ^[77]^, it is feasible that abundant tissue-associated *Endozoicomonas* with GUS activity, including Clade-A isolates, could play a key role in steroid hormone metabolism and possibly in coral broadcast spawning timing and behaviour. Unravelling the connections between host and microbial hormone metabolism pathways will likely reveal new insights into the co-evolution of corals and microbes.

### Potential mechanisms for lipid degradation and uptake from coral hosts or photosymbionts

When in excess, fixed carbon translocated from Symbiodiniaceae is stored as energy reserves within the host. The main storage form is as lipids such as triacylglycerol (TAGs), wax esters, phospholipids and free fatty acids (FAs), which roughly account for 10–40% of total dry coral biomass ^[78–82]^. These compounds therefore represent a significant potential energy source for other holobiont members such as the *Endozoicomonas*. As TAGs are large molecules that cannot enter cells directly, they require enzymatic digestion. Clade-A members possess the genomic potential to degrade TAGs into free fatty acids using the extracellular TAG lipase HlyC, accompanied by the essential lipase chaperone LifO required for proper protein folding during transit through the periplasm in the extracellular environment (Fig. 4). Additionally, both clades have the metabolic capacity to acquire long-chain fatty acids from their extracellular environment through the FadL protein, which facilitates diffusion (Fig. 4) ^[83,84]^. The co-occurrence of members from both clades within the same aggregates ^[32]^ highlights the potential for a cooperative relationship, wherein Clade-B could leverage Clade-A’s ability for extracellular breakdown of TAGs, enabling the utilisation of free fatty acids for various metabolic processes. It is known from model systems such as *E. coli* that long-chain fatty acids can be metabolised, however, carbohydrates are the preferred carbon sources for bacteria when available[85,86].

**Figure 4.**
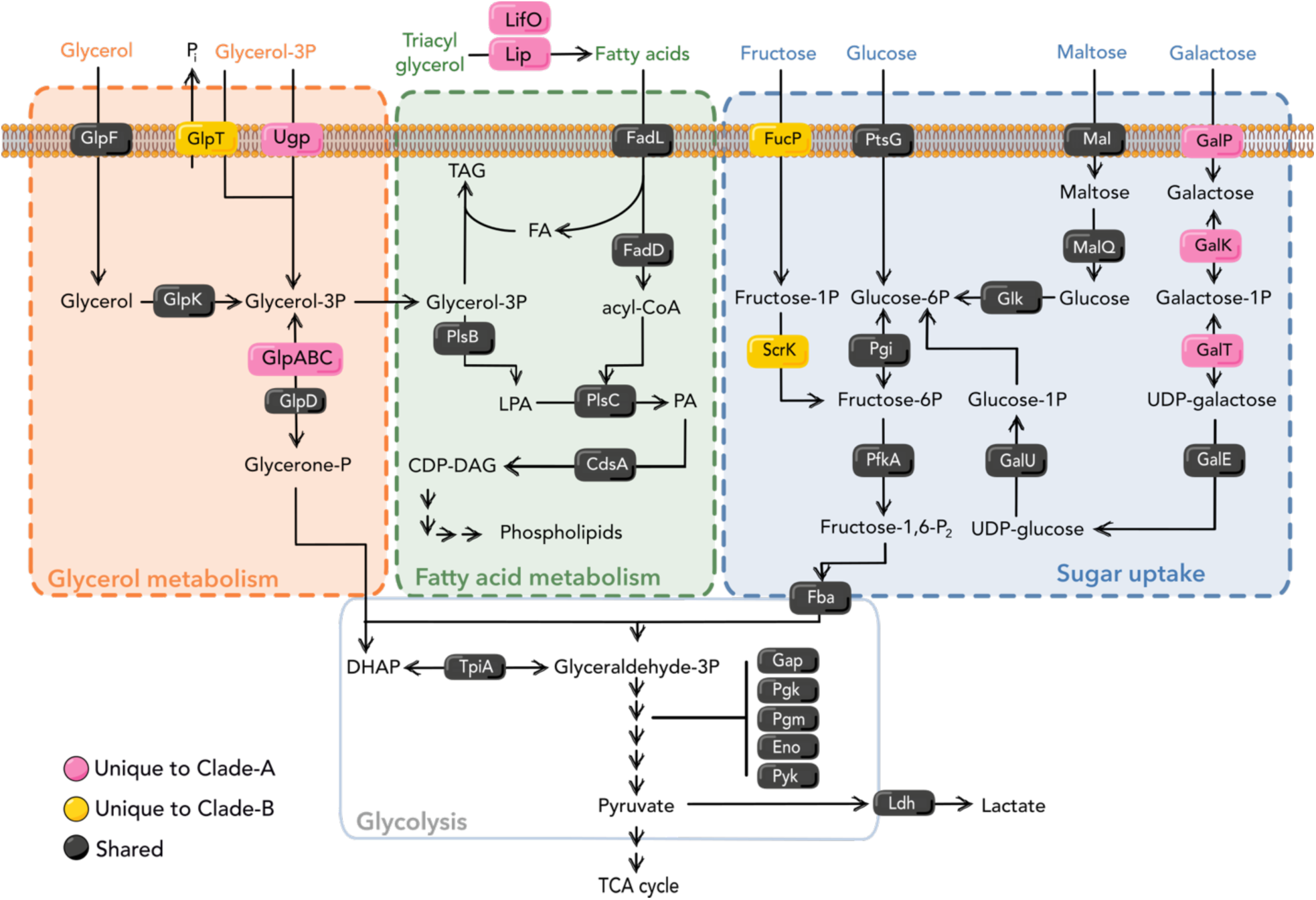
Proposed mechanisms of photosynthate utilisation in *A. loripes*-associated *Endozoicomonas*. General overview of carbon source uptake and degradation pathways within *Endozoicomonas* Clade-A and B based on genomic data. Proteins located on the membrane represent transporters into the bacterial cytoplasm. Pathways with multiple arrows represent multiple reactions not of focus in this study. P, phosphate; TAG, triacylglycerol; DAG, diacylglycerol; CoA, coenzyme; CDP, cytidine diphosphate; LPA; lysophosphatic acid; PA, phosphatic acid; UDP, uridine diphosphate; DHAP, dihydroxyacetone phosphate; Glyceraldehyde-3P, glyceraldehyde-3-phosphate. Magenta: proteins unique to Clade-A; Yellow: proteins unique to Clade-B; Black: shared proteins.

The extracellular hydrolysis of triacylglycerols (TAGs) yields free fatty acids (FAs) and glycerol. Glycerol can be taken up by bacteria via the glycerol facilitator GlpF or phosphorylated intracellularly by glycerol kinase (GlpK) to form glycerol-3-phosphate (G3P). The UgpBAEC (Ugp) transporter system imports both glycerophosphodiesters and G3P, with glycerophosphodiesters hydrolyzed into G3P and alcohol during transport ^[87]^. While the Ugp system is present only in Clade-A, both clades possess GlpF, facilitating glycerol uptake. In Clade-B, G3P can also be imported through the GlpT transporter, coupled with the export of inorganic phosphate (Pi) ^[88,89]^.

After G3P is transported into the cytoplasm, it can either be incorporated into phospholipids ^[90,91]^ or, depending on the availability of (alternative) electron acceptors, it can enter the glycolysis via glyceraldehyde-3-phosphate, thereby enabling the glycolytic catabolism of glycerol (Fig. 4) ^[92]^. Under aerobic conditions, both clades can use G3P for growth via the G3P dehydrogenase GlpD. In contrast, only Clade-A can use fumarate as a terminal electron acceptor for anaerobic growth via the G3P dehydrogenase GlpABC (Fig. 4) ^[93]^.

These observed differences in gene content between the two clades suggest that *Endozoicomonas* has evolved distinct strategies for glycerol metabolism and transport, potentially using glycerol as a major carbon source. Glycerol may be derived directly from photosynthates or through the breakdown of host lipids by TAG lipase. While direct uptake of host-derived glycerol by *Endozoicomonas* has yet to be experimentally demonstrated, certain *Endozoicomonas* strains have been shown to utilize glycerol in liquid culture ^[94]^, and the metabolic potential for anaerobic glycerol degradation has been observed in other species in addition to Clade-A ^[17]^.

#### The potential role of *Endozoicomonas* in phosphorus acquisition

Coral-associated microbes must compete for phosphorus (P) in both organic (P_o_) and inorganic (P_i_) forms within the animal host. Due to the essential roles of P as a structural component of membranes and nucleic acids, bacteria have evolved adaptive strategies to address P_i_ limitations via the utilisation of biologically available dissolved organic phosphorus (DOP). However, unlike P_i_, DOP cannot be directly assimilated by bacteria. Instead, they have evolved multiple transporters and scavenging enzymes such as alkaline phosphatases, phosphodiesterases and nucleotidases to facilitate the utilisation of organophosphates.

Alkaline phosphatases are widely found in marine bacteria where they facilitate the breakdown of phosphoesters (C-O-P bonds) and phosphonates (C-P bonds) in P_o_ compounds to liberate soluble phosphate ^[95–97]^. In this study, we identified two putative alkaline phosphatases (APases): *phoX*, found in both clades, and *phoD*, found only in Clade-B (Fig. 5). Both proteins contain a signal peptide and are therefore expected to be extracellular. Once released, phosphate bound to carrier proteins can traverse the inner membrane via the high-affinity phosphate transport (Pst) system. Smaller organophosphate compounds may also be transported intact across the membrane into the cytoplasm, as hydrophilic molecules <600 Da can pass through porins in the outer membrane into the periplasm ^[98,99]^. Here they can be further translocated into the cell via the G3P uptake system (Ugp) (Figs. 3 & 4) ^[96]^; however, this was only detected in Clade-A. Once internalised, phosphate moieties can be cleaved via a glycerophosphodiester phosphodiesterase GlpQ present in both clades, thereby providing a source of P_i_ but also carbon and nitrogen sources (Fig. 5).

**Figure 5.**
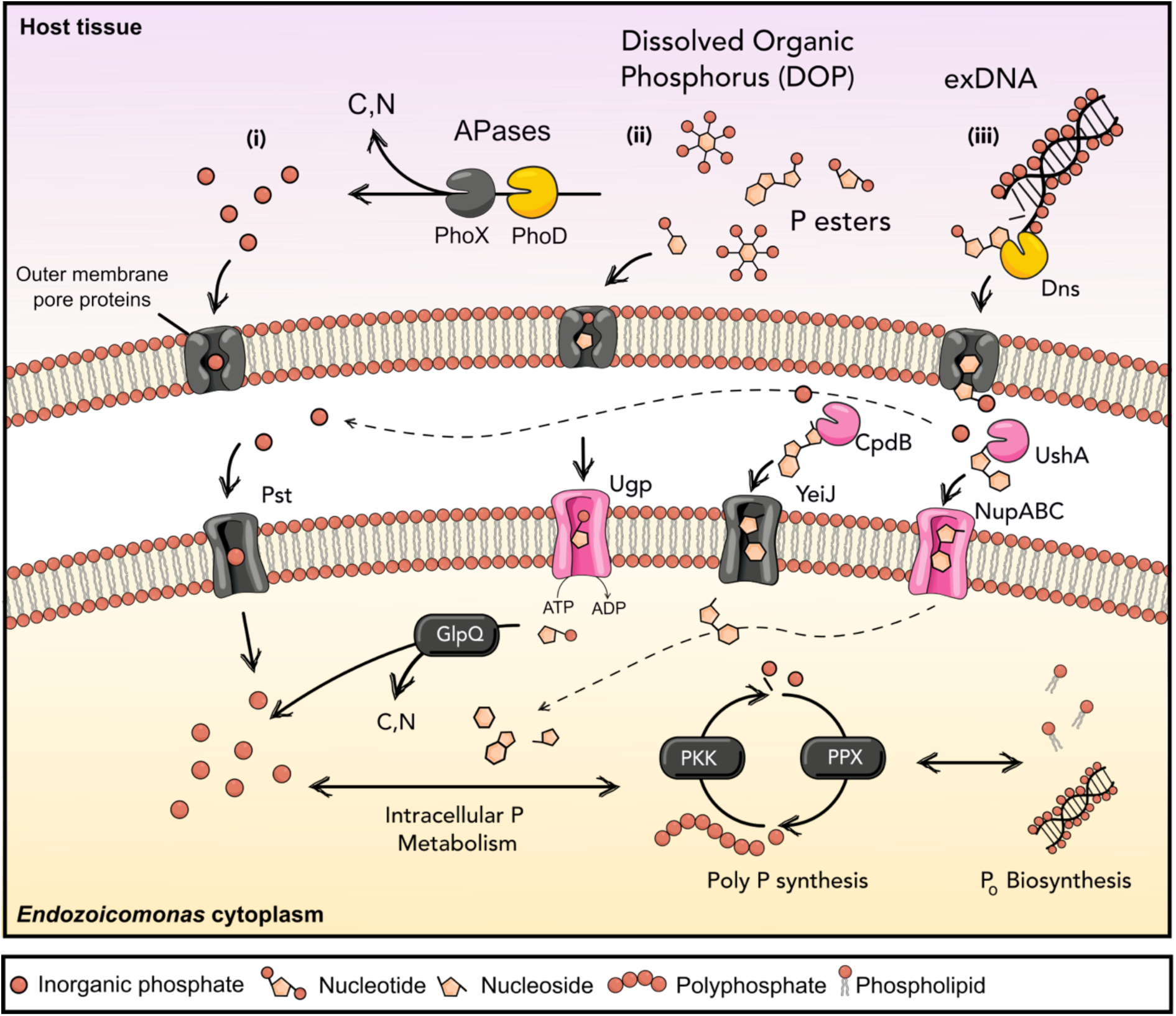
Proposed mechanisms for phosphate assimilation in *A. loripes*-associated *Endozoicomonas*. (i) Extracellular enzymes (predominately APases) result in the release of excess inorganic P (P_i_) followed by the assimilation into the cell. The cleaved organic moiety is either released or taken up via alternative uptake mechanisms and used as carbon/nitrogen sources. (ii) Transport of small P-linked esters via the Ugp-encoded ABC transport system, followed by hydrolysis via the cytoplasmic glycerophosphodiester phosphodiesterase GlpQ. (iii) Extracellular DNA is broken down by the endonuclease Dns, by cleavage at the 3′ carbon, leaving a 5′ phosphate attached to the DNA strand. Once produced, nucleotides can pass across the outer membrane into the periplasm through porins. Periplasmic UshA and CpdB can then remove phosphate groups from 5′ and 3′nucleotides respectively. Inorganic phosphate groups can then traverse the inner membrane via the Pst/PhoU systems, whereas nucleosides can pass through nucleoside transporters (Nup). Magenta: proteins unique to Clade-A; Yellow: proteins unique to Clade-B; Black: shared. Illustration inspired by P metabolism and DOP cycling from ^[24,96,100–102]^.

Extracellular DNA (exDNA) and nucleic acids present in the aquatic milieu represent another major source of organophosphate, but remain unquantified within the holobiont ^[103]^. *Endozoicomonas* clades in this study form CAMAs within their native coral host ^[32]^. exDNA may be readily available within coral tissues or in densely packed microbial aggregates ^[18,104]^. exDNA is found as an integral component of the biofilm extracellular polymeric substances, where it interacts with other EPS components to form a cohesive network required for biofilm stability ^[105–107]^. The utilisation of exDNA as a phosphate source is dependent on secreted endo- and exonucleases to cleave it into nucleotides. *Endozoicomonas* Clade-A genomes encode two periplasmic 5’ and 3’ nucleotidases that are homologous to UshA and CpdB nucleases of the aquatic bacterium *Vibrio cholerae* (Fig. 5) ^[102]^. The UshA and CpdB homologs found in Clade-A shared 50% and 62% amino acid similarity, respectively, with those found in *V. cholerae*. Moreover, Clade-B genomes encode an extracellular deoxyribonuclease *dns* that shares 60.8% protein homology with a nuclease (Dns) from *V. cholerae* ^[104]^. If these nucleases have homologous functions in *Endozoicomonas*, they can allow the use of extracellular DNA as a source of phosphate and nutrition by dephosphorylating extracellular nucleotides before they are internalised within the cell ^[102]^. This is corroborated by the finding of the transporter NupABC in Clade-A responsible for nucleoside uptake ^[108]^ and the homologous nucleoside transporter YeiJ found in both clades (Fig. 5) ^[109]^. Despite differences in gene content between clades, they may have evolved to fulfil similar roles. Moreover, multispecies aggregates containing members from both clades ^[32]^ point to the potential of different species with different gene content partially contributing to the same phosphorus-scavenging pathway, i.e., via metabolic complementation ^[51]^.

Both the coral host ^[110–112]^ and Symbiodiniaceae ^[113,114]^ have been found to express APases, however, the significance of bacterial APases within the coral holobiont remains uncertain. Nonetheless, microbial P_i_ transporters and enzymes involved in the uptake and degradation of organic molecules containing P are upregulated in other marine bacteria in a *pho-*dependent manner when P is scarce ^[115]^. This includes phosphate starvation genes such as *dns, ushA* and *phoX, phoD* ^[116]^. Moreover, CpdB activity is also induced under phosphate-limiting conditions, however the molecular mechanisms are unknown ^[102]^. The pho regulon is controlled by a two-component regulatory system, composed of a transmembrane histidine kinase (PhoR) and a response regulator (PhoB) ^[117]^, both of which are present in the genomes of each clade. This suggests a possible functional role in P scavenging within the holobiont during P limitation.

An analogy to another symbiosis is the expression of APases by soil microorganisms, which plays a vital role in soil P_o_ remineralisation by enhancing extracellular P_i_ availability for host plant roots ^[118,119]^. Likewise, tissue-associated *Endozoicomonas* expressing extracellular APases that release P_i_ could provide readily available P for the entire holobiont, benefiting other microbial members near *Endozoicomonas* unable to directly access organophosphates, i.e., Symbiodiniaceae. Moreover, it was observed, that members of Clade-B encoded an organophosphate: the P_i_ antiporter family transporter (GlpT), which facilitates the transport of P_O_ across biological membranes with the simultaneous translocation of inorganic phosphate in the opposite direction (Fig. 5) ^[89]^. Depending on glycerol concentration in the extracellular milieu, this may also deliver P_i_ to Symbiodiniaceae.

Conversely, the upregulation of microbial scavenging enzymes and high-affinity transport mechanisms during P limitation could potentially have adverse effects on holobiont functioning. A transition from limited to excess nitrogen can induce P starvation in Symbiodiniaceae as the excess N stimulates growth ^[120]^. This can occur during thermal stress, when host catabolism increases, resulting in an increased availability of ammonium ^[121]^. P limitation affects the lipid composition and functioning of the Symbiodiniaceae chloroplast, as phospholipids in the thylakoid membranes are substituted by alternate lipids, e.g., sulfolipids ^[122]^. This substitution compromises membrane stability under thermal stress leading to reactive oxygen species leaking into the host tissues triggering the expulsion of Symbiodiniaceae, a phenomenon known as coral bleaching ^[28,29,31]^. Therefore, if dominant *Endozoicomonas* communities exhibit high P-scavenging abilities *in hospite*, as suggested by their P intracellular enrichment within coral tissues ^[24]^, this could exacerbate P limitation if their scavenging mechanisms surpass those of the coral host or photosymbiont. Hence, understanding how abundant tissue-associated *Endozoicomonas* can influence P availability and cycling within the coral holobiont during stress is crucial for unravelling the cellular mechanisms behind coral bleaching, especially in increasingly warm and nitrogen-enriched coastal waters ^[123,124]^.

#### Underlying mechanisms governing *Endozoicomonas* aggregation patterns

In a previous study, we revealed distinct clade-specific aggregation morphologies co-occurring with the same coral host ^[32]^. In CAMAs formed by Clade-A, bacteria exhibited aggregates characterised by well-defined boundaries and contained growth patterns, while Clade-B members formed clusters lacking clear boundaries and displaying unrestricted growth ^[32]^. This stark contrast prompted our investigation into identifying underlying genomic signatures potentially driving these differing aggregation patterns and their potential implications for microbial interactions within the coral host.

Molecules on the outermost surface layer of bacteria play essential roles in bacteria-host interactions. They allow hosts to recognise pathogens and in turn, mount an immune response, and they also play a vital role in mutualistic animal-microbe interactions, as demonstrated in the well-described legume-rhizobia symbiosis ^[125,126]^. Outer capsular polysaccharides (CPSs) are complex carbohydrate structures considered to act as protective barriers around bacterial cells ^[127]^. They provide several advantages, such as shielding Microbe-Associated Molecular Patterns (MAMPs) from recognition by host immune defences facilitating immune cell evasion and adhesion to host cells, thereby assisting with tissue colonisation ^[128–130]^. Moreover, CPSs are also considered determinants of biofilm physicochemical properties and have been demonstrated to cause auto-aggregation of cells ^[131–133]^. Yet, adhesion, biofilm formation, and their roles in host colonisation and persistence remain understudied in *Endozoicomonas*.

Several complete pathways for the synthesis of CPSs that were identified in members of Clade-B are lacking in Clade-A. These include the biosynthesis of UDP-N-acetyl-mannosamine (UDP-ManNAc), GDP-L-fucose, (GDP-L-Fuc) and CMP-N-acetylneuraminate (Neu5Ac/sialic acid) (Fig. S5). Capsules composed of sialic acid have garnered much interest in clinical microbiology as they can help bacteria evade the host immune system by mimicking eukaryotic cell surface glycoconjugates, thereby hindering the recognition of invading bacteria by immune cells ^[134–136]^. While sialic acids are poorly characterised in scleractinian corals, evolutionarily conserved sialic acid receptors have been reported on the cell wall of Symbiodiniaceae ^[137]^. Sialic acid capsules could represent a way for Clade-B *Endozoicomonas* bacteria to evade the host phagocytosis by mimicking the cell surface of photosymbionts, and this may therefore explain their unrestricted growth ^[138,139]^. Notably, the complete pathway required for the synthesis of sialic acid was only found in four strains of Clade-B, which include the by far most dominant strains ALB032, ALC066 (combined relative abundance of ∼ 50% ^[32]^).

Conversely, Clade-A contains an LPS gene cluster homologous to the Wbp pathway in *Pseudomonas aeruginosa* (Fig. S5), which encodes all five enzymes required for the *de novo* biosynthesis of the uronic acid monosaccharide di-N-acetylated mannosaminuronic acid typically found in the outermost portion of LPS of the Gram-negative cell wall (O-antigen layer) ^[140,141]^ (Fig. S5). The O-antigen layer is highly variable, differing at both species and strain levels. It has been observed in both pathogenic and non-pathogenic Gram-negative bacteria but is best described in the respiratory pathogenic bacteria *Bordetella pertussis* and *Pseudomonas aeruginosa*, where it confers resistance to multiple host defences ^[142–144]^. Uronic acids, like mannuronic and glucuronic acid, constitute part of the capsule-like exopolysaccharide (EPS) matrix in *Pseudomonas aeruginosa*, a model organism for biofilm research ^[145]^. These acids confer anionic properties to the biofilm, facilitating the binding of divalent cations such as calcium and magnesium ^[146,147]^, resulting in a more robust and ‘sticky’ matrix ^[148]^. Thus, the potential scavenging and conversion of host uronic acids via GUS enzymatic activity in Clade-A may not only provide a carbon source but could potentially also serve as building blocks for EPS formation. Notably, the metabolic capacity to synthesise UDP-ManNAc and CMP-N-ac and sialic acid were exclusive to strains in Clade-B (Fig. S5). This suggests divergent cell-cell attachment strategies within phylogenetically distinct *Endozoicomonas*, possibly contributing to the different morphology in CAMAs formed by Clade-A and Clade-B.

While invasion strategies are best characterised in pathogenic bacteria, it is plausible that mutualistic bacteria have evolved analogous mechanisms to establish associations within their respective hosts. Thus, the molecular architecture of the bacterial cell surface represents an ecologically relevant avenue for studying bacterial interactions with host proteins and their role in colonisation ^[149–151]^. Considering this, genomic differences in the suite of bacterial extracellular polysaccharides may explain the differences in the CAMA aggregation pattern between Clade-A and Clade-B that was observed *in hospite* ^[32]^.

### Factors facilitating evasion of host immune responses

Genomes in both clades encode a high number of eukaryotic-like proteins (ELPs), e.g., WD40, ankyrin and Sel1-like repeat sequences, with Clade-B exhibiting notably higher counts of WD40 and ankyrin repeats compared to Clade-A, whereas Sel1-like repeats were exclusively enriched in Clade-A (Table S3). ELPs have previously been identified in a number of other *Endozoicomonas* genomes where they are hypothesised to be involved in host tissue internalisation and persistence via protein-protein interactions ^[16,21,25,26]^. This is supported by the observed upregulation of ankyrin-containing proteins in *E. marisrubri* 6c upon exposure to host tissue ^[27]^. Of particular interest in this study was the ankyrin-containing protein AnkX which encodes a phosphocholine (ChoP) transferase. This was found in five copies in one Clade-B strain (AL032), which dominated the *in hospite* bacterial communities of *A. loripes* ^[32]^. In the intracellular human pathogen *Legionella pneumophila*, AnkX is employed to modify the eukaryotic host membrane trafficking proteins, suppressing phagocytosis and increasing bacterial survival within host cells ^[152]^. Notably, Clade-B was found to encode an extracellular phospholipase C (PLC) capable of cleaving ChoP headgroups from the major phospholipid ^[153]^, particularly phosphatidylcholine, which are major components of cell membranes ^[153]^. By cleavage of ChoP, PLC can disrupt the integrity of host cell membranes, aiding in bacterial invasion. There is some evidence, that both commensal and pathogenic bacteria in humans utilise ChoP cleaved from host membranes to modify their bacterial surface or secreted products thereby reducing the immune response to colonisation ^[154,155]^. This suggests that members of Clade-B, which were found to be the dominant *Endozoicomonas* lineage within *A. loripes*, may have the functional potential to utilise host-derived ChoP to modulate the host immune responses.

### Secreted Factors Facilitating Host Colonisation and Competition

The *Endozoicomonas* genomes possessed a genetic repertoire related to host colonisation and adhesion, with multiple near-complete secretion systems identified, including type I, II, III, and VI. While almost all secretion systems were detected in both clades, a near complete type VI secretion system (T6SS) gene cluster encoding a T6SS apparatus was solely identified in Clade-A (Table S4 and Fig. S6). The T6SS was positioned within a gene cluster on a single contig of strain ALB091, ALB115, ALE010 determined by Kyoto Encyclopedia of Genes and Genomes (KEGG) orthology and conserved domain search (Fig. S7). The T6SS of gram negative bacteria is known to play an important role in inter-bacterial competition via the delivery of toxic effector molecules into adjacent cells ^[156,157]^. Additionally, recent studies have found that effectors targeting the eukaryotic host can also be utilised to facilitate host invasion or gain access to intracellular host resources, ultimately assisting the attacking bacteria in acquiring a competitive fitness advantage ^[158]^. The diversity of these translocated molecules referred to as Type VI secretion system effectors (T6SSE), can serve a range of functions within target cells depending on the bacterial species ^[159,160]^. Therefore, investigating potential T6SSEs in *Endozoicomonas* and how they exert their action upon recipient cells could enhance our understanding of how they can outcompete other bacteria *in hospite* as well as provide insight into the establishment and maintenance of the host-microbe relationship.

Proteins predicted to be T6SSE included an A1/A2 phospholipase (PLA), and the alkaline phosphatase PhoX (Table. S5). PLA is an outer membrane lipase, previously described in *Pseudomonas* spp. and *E. coli* to hydrolyse ester bonds of both phosphatidylserine (PS) and phosphatidylcholine (PC) ^[161–163]^. The outer membrane of Gram-negative bacteria such as *E. coli* consists of up to 70% phosphatidylethanolamine (PE), with very little PS and PC in their membranes. While not much is known about the phospholipid composition of *Endozoicomonas*, the predominant polar lipids of three *Endozoicomonas* type strains consist of PE, phosphatidylglycerol (PG), and diphosphatidylglycerol ^[35,164,165]^. Therefore, PLA might help coral-associated *Endozoicomonas* outcompete other bacterial communities with PC and PS as the main membrane phospholipids (Fig. 6B)^[163]^. PLA could also be employed to trigger internalisation within host cells by disruption of host membranes (Fig. 6A) ^[166]^, as phospholipids in a range of coral species are comprised of PC and PS ^[167]^. Consequently, the contained aggregation pattern formed by members of Clade-A could be due to an intracellular location contained within a host-derived membrane ^[32]^. Conversely, *Endozoicomonas* Clade-B members lack T6SS and have unrestricted growth patterns suggesting an extracellular location within the host tissues. Additionally, a microbial collagenase, ColA was also predicted as a T6SSE, which could further aid in host infection, as collagenous fibres are major components of the extracellular matrix of corals (mesoglea) ^[168,169]^.

**Figure 6.**
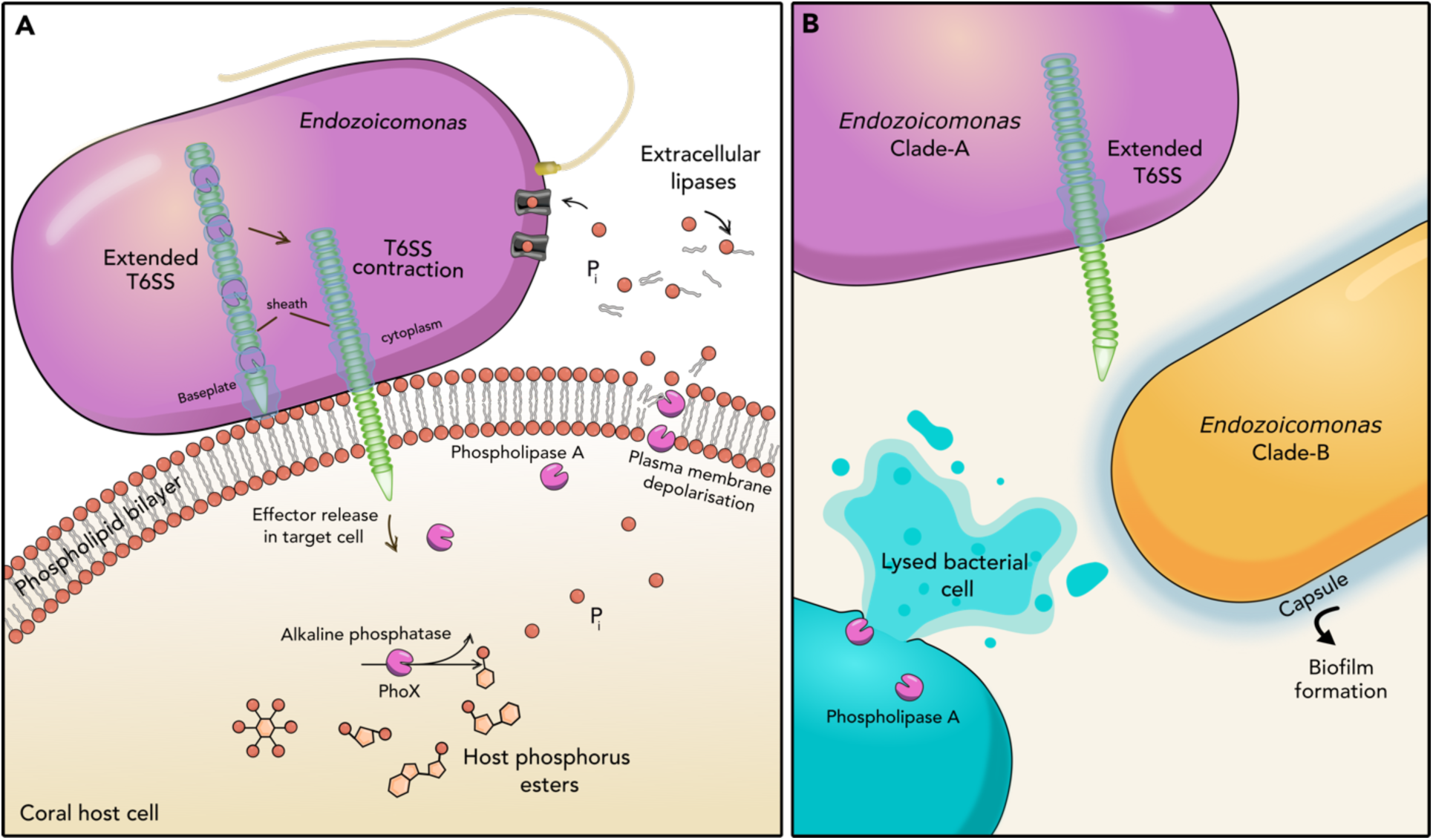
Type VI secretion system-mediated delivery of *Endozoicomonas* effector proteins. (A) Model of effector protein delivery by the Clade-A Type VI secretion system (T6SS) and the potential impact of these effector proteins on the host. The T6SS consists of a sheath (green) which contracts to inject a cell-puncturing structure into a neighbouring cell, releasing effector proteins. Effector proteins are either bound within the lumen of the spike (A), or on the exterior (not depicted). For instance, phospholipase A may induce plasma membrane depolarisation, potentially facilitating host invasion through a puncture hole, while alkaline phosphatase may influence phosphorous cycling. (B) Hypothesised interactions involving *Endozoicomonas* and other coral-associated bacteria. Phospholipase A may target phosphatidylcholine and phosphatidylserine in competing bacterial membranes (blue cell), giving Endozoicomonas Clade-A an advantage. However, the coexistence of Clade-A and Clade-B within the same aggregate ^[32]^ suggests non-antagonism, potentially due to capsule formation in Clade-B which could prevent direct cell-to-cell contacts required for translocation of effector proteins ^[170]^.

While lipases may facilitate the colonisation of host tissues via the disruption of host cell membranes, they can also play a role in nutrient acquisition. T6SS-delivered phospholipase effectors with PLA activity have the functional potential to act on host cell membranes (releasing choline, P_i_), as well as storage lipids, as PC and PS are also part of the composition of many storage lipids ^[167]^. This is corroborated by the finding that another predicted T6SSE, phoX, could be translocated into host cells to liberate phosphate from organic phosphorous compounds within the host or in prokaryotic competitors.

The T6SS was found in roughly half of published *Endozoicomonas* genomes (Fig. 7), exhibiting a sporadic distribution pattern across phylogenetic lineages, possibly suggesting multiple potential horizontal gene transfer events. In contrast, PLA appears to be a common feature in *Endozoicomonas* and was detected in all genomes used in this study except strain ALC013 from Clade-A, which was the least abundant strain within the *A. loripes* microbiome (>0.05% relative abundance). The presence of PLA in isolates without a T6SS indicates that these lipases likely also serve alternative functions beyond host colonisation and bacterial competition. Indeed, PLAs are responsible for fast turnover rates of cellular phospholipids, playing central roles in membrane maintenance and remodelling ^[174]^.

**Figure 7.**
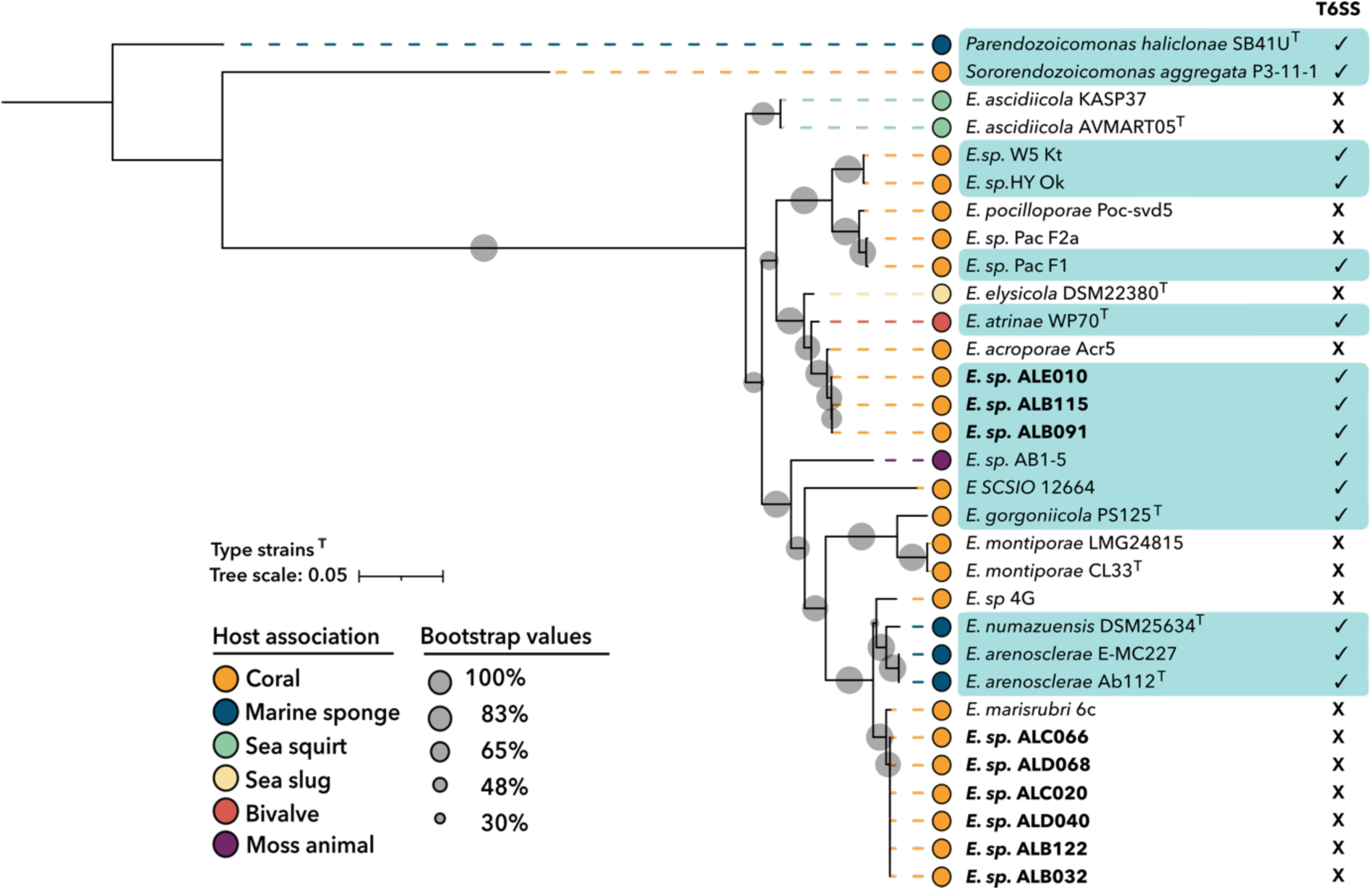
Phospholipase A1 phylogeny of *Endozoicomonas* genus, based on published genomes. Phospholipase A1 protein sequences were aligned using MAFFT v 7.487 ^[171]^. The ML tree was constructed with IQ-tree2 v1.6.1 ^[172]^ using the LG+G4 model selected by ModelFinder Plus. The final tree was visualised in iTOL v6 ^[173]^ and rooted using *Parendozoicomonas haliclonae* S-B4-1 as the outgroup. Strain colours correspond to the source organism, and T6SS indicates Type VI secretion system presence in the respective genomes, which are highlighted in teal.

Notably, among the two *Endozoicomonas* clades analysed, only members of Clade-A possessed the genomic repertoire encoding a T6SS. Bacteria have evolved various mechanisms to compete and survive in complex microbial communities. Capsule formation is one such strategy that in some bacteria can serve as a protective layer against antagonistic factors such as (T6SS) present in certain microbes ^[175–178]^. This could perhaps explain why Clade-A and Clade-B can co-exist within the same host, as well as within the same CAMA ^[32]^ if the predicted capsule layer of Clade-B is expressed as suggested by Clade-specific differences observed *in vitro* (Fig. S8). Conversely, this could also be a case of bacteria having ‘immunity’ towards T6SSE secreted from closely related bacteria ^[179,180]^.

## Conclusions

Our findings underscore the metabolic adaptability of *Endozoicomonas* bacteria and their potential contributions to coral holobiont dynamics. Metabolic reconstruction revealed previously undescribed enzymatic repertoires, indicative of the utilisation of complex host-derived compounds, including starch, lipid compounds, and the scavenging of organic phosphorus and hormone-conjugates (Table 2). Moreover, previously undescribed genomic features were identified, suggesting diverse host colonisation strategies within each clade, either through modifications to the outer surface structures or secretion of directed T6SSE. The widespread occurrence and abundance of *Endozoicomonas* in healthy corals suggests a reciprocal resource exchange, where the number of resources provided depends on the amount of resources received. Future studies should aim to investigate putative beneficial functions of nutrient cycling involving simple sugars and inorganic phosphorus, and their potential impacts on host and/or photosymbiont dynamics in homeostasis, as well as under heat-induced nutrient limitation. This may include examining how *Endozoicomonas* responds to phosphate starvation by assessing the expression of genes involved in phosphorus assimilation, including those encoding inorganic phosphate and glycerol transporters, extracellular phosphatases, nucleases and esterases.

**Table 2.**
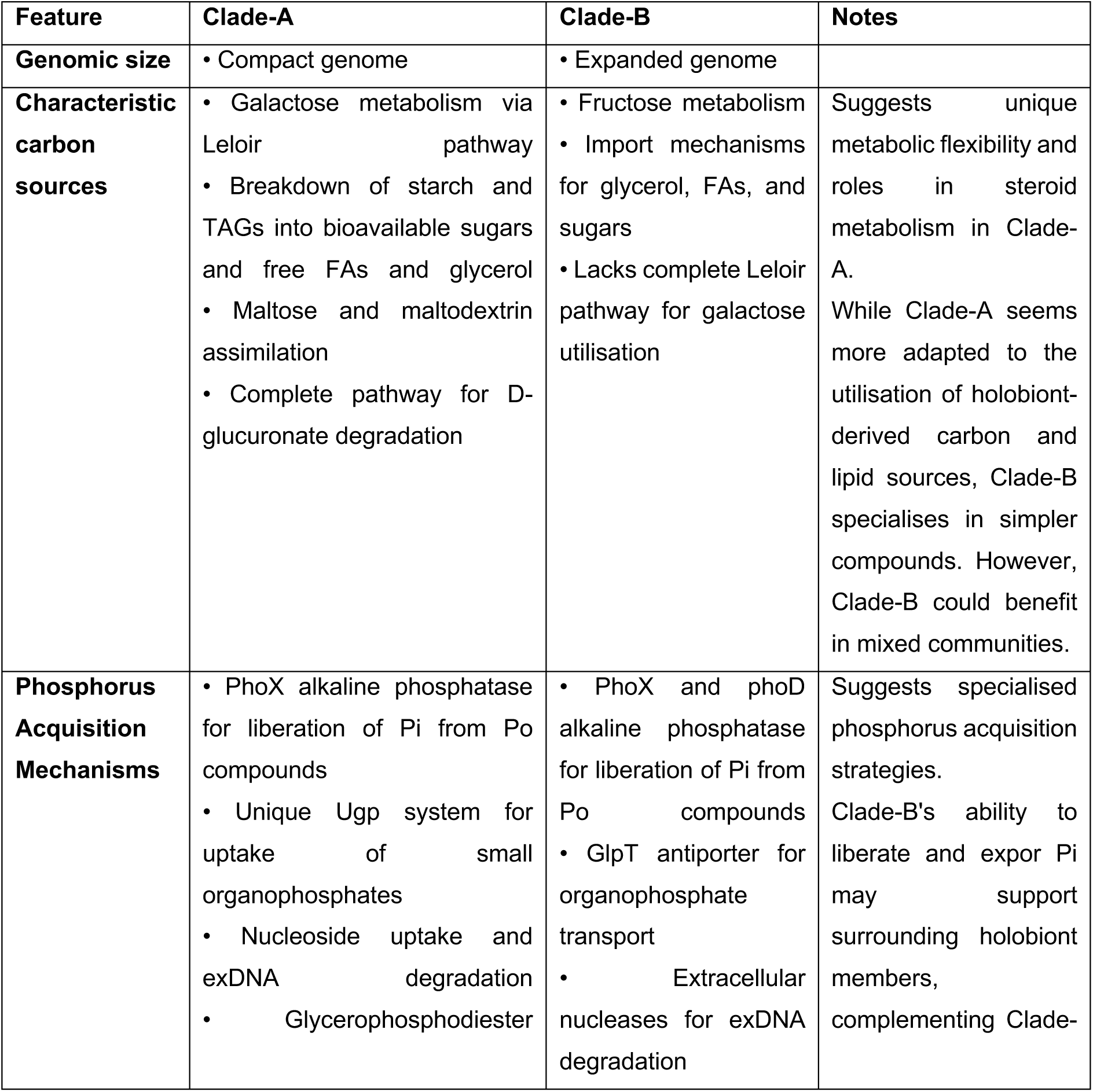
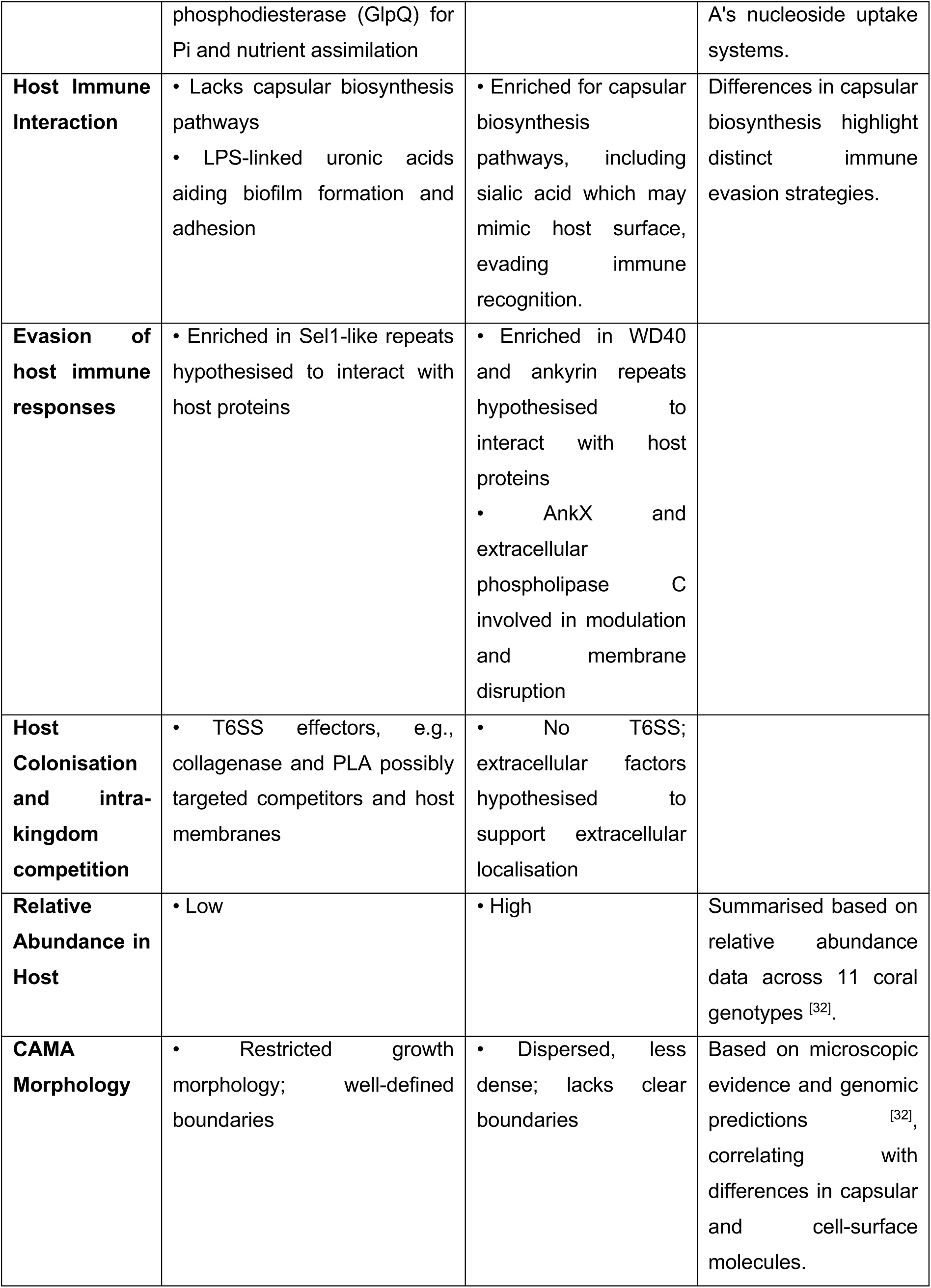
Summary of Main Differences Between *Endozoicomonas* Clade-A and Clade-B.

## Materials and Methods

### Isolation of *Endozoicomonas* cultures and their genomic DNA

The methods used to isolate and culture *Endozoicomonas* were previously described in ^[32]^. Briefly, the *Endozoicomonas* 11 strains (Table 1) used in this study were isolated from the stony coral *A. loripes* (Acroporidae) collected from two reef sites in the central Great Barrier Reef in Australia; Davies Reef (18°51’S, 147°63’E) and Backnumbers Reef (18°31’23.4“S 147°08’39.9”E). Approximately 3 cm-sized coral fragments were washed using filter-sterilised seawater (SSW) to remove loosely associated microbes and other debris. Each coral fragment was tissue blasted into 5 mL of filtered seawater (pore size 0.2 μm). The resulting tissue slurry was homogenised using a tissue homogeniser (HoverLabs, India) to break down tissue clumps and immediately plated in triplicate onto Marine Agar 2216 medium (BD Difco) and incubated at 23 °C in the dark. Single colonies were obtained after a minimum of three clean passages onto fresh plates.

A volume of 5 mL of marine broth (2216 BD Difco) (MB) was inoculated with one bacterial colony and kept under continuous shaking (175 rpm) at 28 °C. After two days of incubation, the bacterial cultures reached the stationary phase and genomic DNA was extracted using two separate protocols, one for short-read Illumina sequencing: Invitrogen™ PureLink™ Genomic DNA Mini Kit (K182001, Invitrogen™, Australia) and one for long-read Nanopore sequencing: Qiagen MagAttract High Molecular Weight DNA Kit (#67563, Qiagen™, Australia). Both protocols were carried out according to the manufacturer’s instructions; however, DNA extracted using the Qiagen™ MagAttract HMW DNA Kit included a 90 min digestion with Proteinase K at 56 °C. The quantity of extracted DNA was measured using the Quant-iT™ PicoGreen™ dsDNA Assay Kit (P7589, Invitrogen™, Thermo Fisher Scientific, Australia) and the quality was assessed using a UV5 Nanospectrophotometer (Mettler Toledo, Australia).

### Whole-genome sequencing and assembly

High-quality genomic DNA was subjected to two different sequencing approaches: the Illumina NextSeq550 platform creating 150 bp paired-end reads and the Nanopore MinION creating long-read reads (Oxford Nanopore), both of which were conducted at Doherty Applied Microbial Genomics (Peter Doherty Institute, Melbourne, Australia). In cases where the depth of the long-read data was sufficient (100x coverage), long-read data was quality checked using Filtlong v2.9.5 (GitHub repository, https://github.com/rrwick/Filtlong) using the parameters min-length 1kb, keep_percent 5. Conversely, if the depth of the long-read data set was insufficient for a long-read-first assembly, only a short-read assembly was performed. Long-read genome assemblies were performed with Flye v.2.9.1 ^[181]^ with default parameters, followed by long-read polishing with Medaka v1.12.1 (Github Repository https://github.com/nanoporetech/medaka) and short-read polishing with high quality, trimmed paired-end Illumina reads with a phred score of 30 using POLCA v4.0.3 ^[182]^. Short Illumina reads were quality-checked using FastQC (Github repository https://qubeshub.org/resources/fastqc), followed by adaptor removal and read trimming using cutadapt v4.8 (phre>30) ^[183]^. Short-read only assembly was carried out with quality-filtered and trimmed reads using SPAdes De Novo Assembler v1.15.5 ^[184]^. Contigs were checked for contamination using CAT/BAT with the Non-Redundant database ^[185]^, and any contigs belonging to a different phylum than the taxonomy assigned by GTDB-Tk was removed. Finally, the assembled genomes were checked for completeness, contamination and strain heterogeneity using CheckM v1.1.0 with the lineage-specific option ^[186]^.

### Genome annotation and characteristics

All publicly available *Endozoicomonas* reference genomes (n=27) were downloaded from the NCBI Genomes database. Gene prediction and annotation of genomes used in this study (11 isolates and 27 publicly available genomes) were conducted using Prokka v1.14.5 with default settings ^[187]^ and Rapid Annotation using Subsystems Technology (RAST) v2.0 ^[188]^. A list of all reference genomes used in this study alongside their genomic characteristics (size, coding density, contig numbers) as well as host organisms are provided in Table S1. Genes of interest were classified based on KEGG orthologies ^[189]^ using the whole proteome as a query within the web-based BlastKOALA tool against the taxonomic database “Bacteria” v2.2 ^[190]^. Pathway completion was estimated using KEGG-mapper ^[191]^. InterProScan v5.55 was used for protein family (Pfam) annotations to identify eukaryotic-like repeat domains using an e-value cut-off of 1 × 10^−5^ as the filtered value ^[192]^. For the annotation of genes of interest, conserved domains were verified with web-based BLASTp-searches using an *e*-value cut-off of 1 × 10^−5^ ^[193]^. SignalP v6.0 ^[194]^ and PSORTb v3.0.3 ^[195]^ servers were used to predict the presence and location of signal peptide cleavage sites in protein sequences with default parameters. A comprehensive list of all metabolic genes/pathways queried in this study can be found in Table S2. Type VI secretion system effector molecules were predicted using web-based Bastion6, with a cutoff score of ≥0.5 ^[196]^. Bastion6 employs a two-layer Support Vector Machine (SVM)-based ensemble model that integrates a wide range of features extracted from known T6SS effector protein sequences, using both unsupervised and supervised learning to identify potential T6SS effectors accurately.

### Phylogenomic Analyses

Whole genome-based similarities among members of the genus *Endozoicomonas* used in this study were estimated using the average nucleotide identity (ANI), average amino acid identity (AAI) and *in silico* genome– genome distances (GGDs). ANI was calculated using the EzBioCloud web-based ANI calculator ^[197]^. AAI was calculated with CompareM (https://github.com/dparks1134/CompareM) and GGDs were calculated using the DSMZ web-based calculator with formula 2 ^[198]^. All heatmaps were generated using ggplot2 v. 3.4.4 ^[199]^ implemented in R v3.4.1 ^[200]^. Phylogenies were inferred via the alignment of 120 bacterial marker genes identified by the Genome Taxonomy Database (GTDB) ^[186]^ via GTDB-Tk v2.1.0 ^[201]^. A maximum-likelihood phylogenetic tree was constructed in IQ-TREE2 v1.6.1 ^[172]^ with a LG + F + R7 model obtained from automatic model selection through ModelFinder plus ^[202]^ with robustness of phylogenetic placement tested through 1000 ultrafast bootstraps. The distantly related bacterial species *Parendozoicomonas haliclonae S-B4-1* ^[203]^, *Sororendozoicomonas aggregata* Pac-P3-11-1 ^[204]^ and SCSIO12664 ^[205]^ were selected as outgroups. The resulting tree was visualised using iTOL v6 ^[173]^. Genome coding densities denoted on the ML tree were defined as the total genome size divided by the total number of genes (excluding non-coding sequences and pseudogenes).

## Supporting information

Supplementary data overview

## Ethics approval and consent to participate

Not applicable

## Consent for publication

Not applicable

## Availability of data and materials

Sequence data that support the findings of this study have been deposited in the NCBI the repository with the primary accession code PRJNA1200206

## Authors’ contributions

C.R.G, L.H, L.L.B and M.J.H.v.O designed the study. Genomic DNA extractions were carried out by A.D. Methodology was established by K.T, J.M and C.R.G. C.R.G, K.T and G.K.P worked on bioinformatics and analysis. C.R.G created visualisations. C.R.G wrote the first draft with all other authors editing and approving the final manuscript.

## Acknowledgements

We acknowledge the Wulgurukaba and Bindal peoples as the first scientists and traditional custodians of the lands and sea country where this research was conducted. We extend our sincere gratitude to the SeaSim team for their collaboration and support in coral collection and husbandry.

## Supplementary material for

**Table S1.**

List of all Endozoicomonadaceae genomes used in phylogenomic analyses.

**Table S2.**

List of all metabolic genes and pathways interrogated to explain the underlying mechanisms behind differing aggregation patterns between *Endozoicomonas* Clade-A and Clade-B, nutrient transfer and uptake, and how Clade-A and Clade-B respond to environmental challenges, including nutrient deprivation and the host immune response. The + indicates whether the KO was present in all strains of the respective clade, (+) indicates that it was present in all strains within one clade except one.

**Table S3. Eukaryotic-like protein sequences.**

Number of eukaryotic-like protein sequences found in the 11 *Acropora loripes-*sourced strains in comparison to publicly available genomes. Sequences were detected based on an Pfam classification (PF). Cad: Cadherin repeat, Sel1: Sel1-like repeat, TPR: Tetratricopeptide repeat, WD40: WD40 repeat, PQQ: Pyrrolo-quinoline quinone repeat, ANK: Ankyrin repeat

**Table S4.**

List of type VI secretion system (T6SS) core subunits (TssA-TssM) and accessory components. The presence/absence of *Endozoicomonas* Clade-A proteins encoding T6SS based on KEGG Orthology. The Tss (Type VI Secretion) genes follow the T6SS gene nomenclature proposed by ^[198]^. These genes are highly conserved across the genomes of more than 100 bacterial species, all of which encode a T6SS similar to the system ^[199]^.

**Table S5.**

List of predicted type VI secretion system effectors (T6SSEs) in the genomes of *Endozoicomonas* Clade-A predicted bioinformatically using the Bastion6 prediction tool. Only proteins with a confidence score greater than 80 are included in this table.

**Figure S1.**
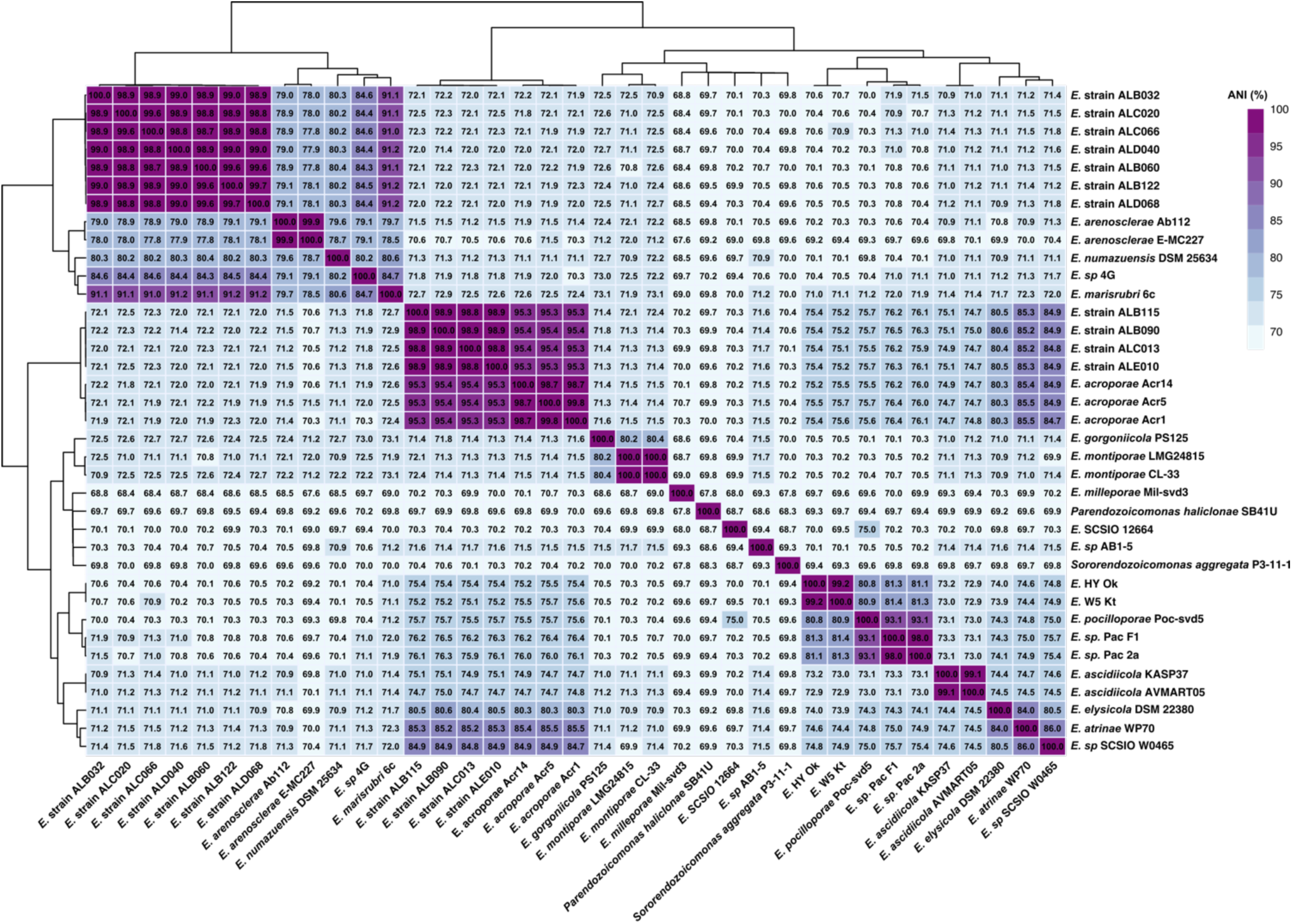
Heatmap showing the pairwise average nucleotide identity (ANI) between isolates obtained in this study and others from the literature.

**Figure S2.**
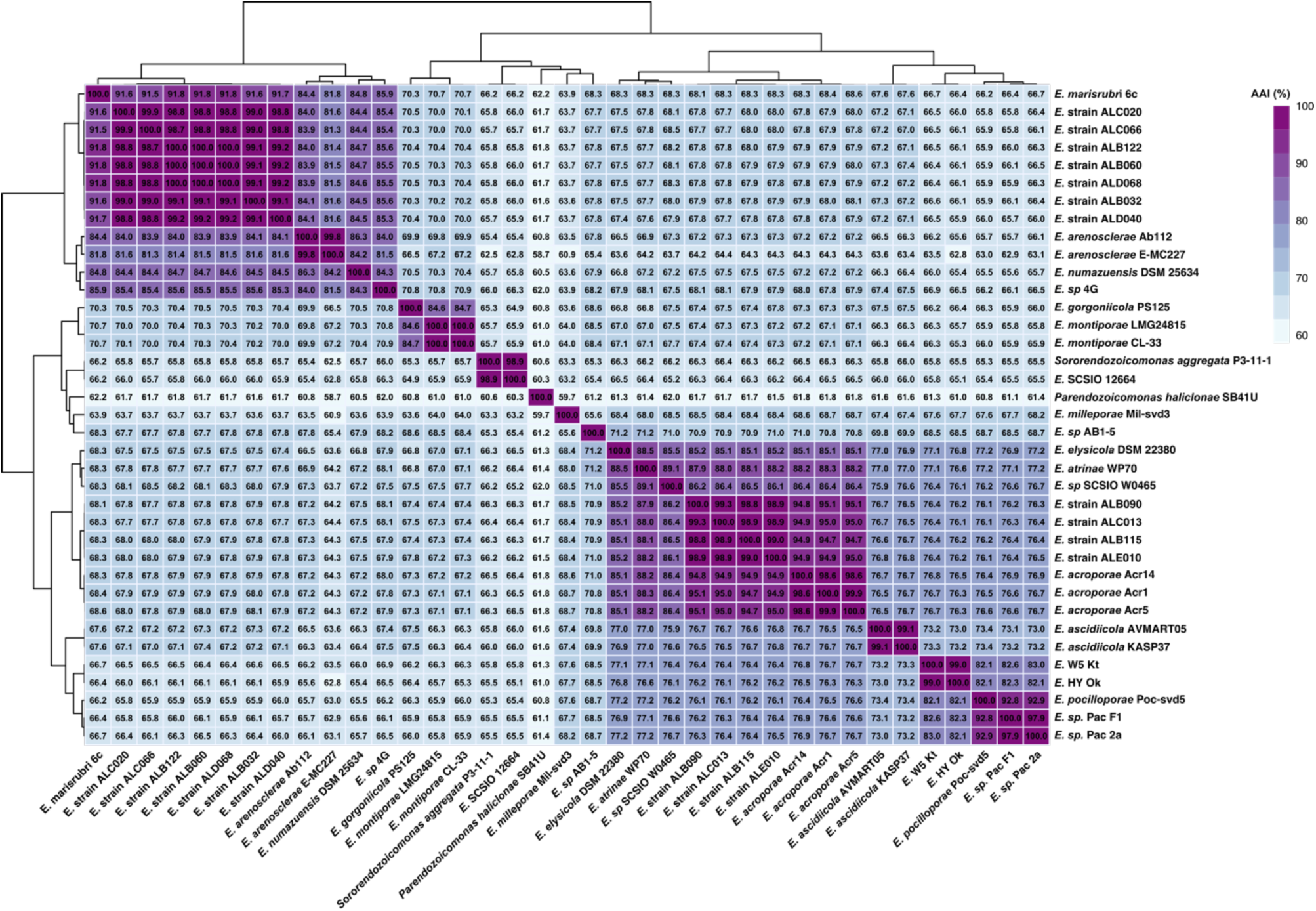
Heatmap showing the average amino acid identity (AAI) between isolates obtained in this study and others from the literature.

**Figure S3.**
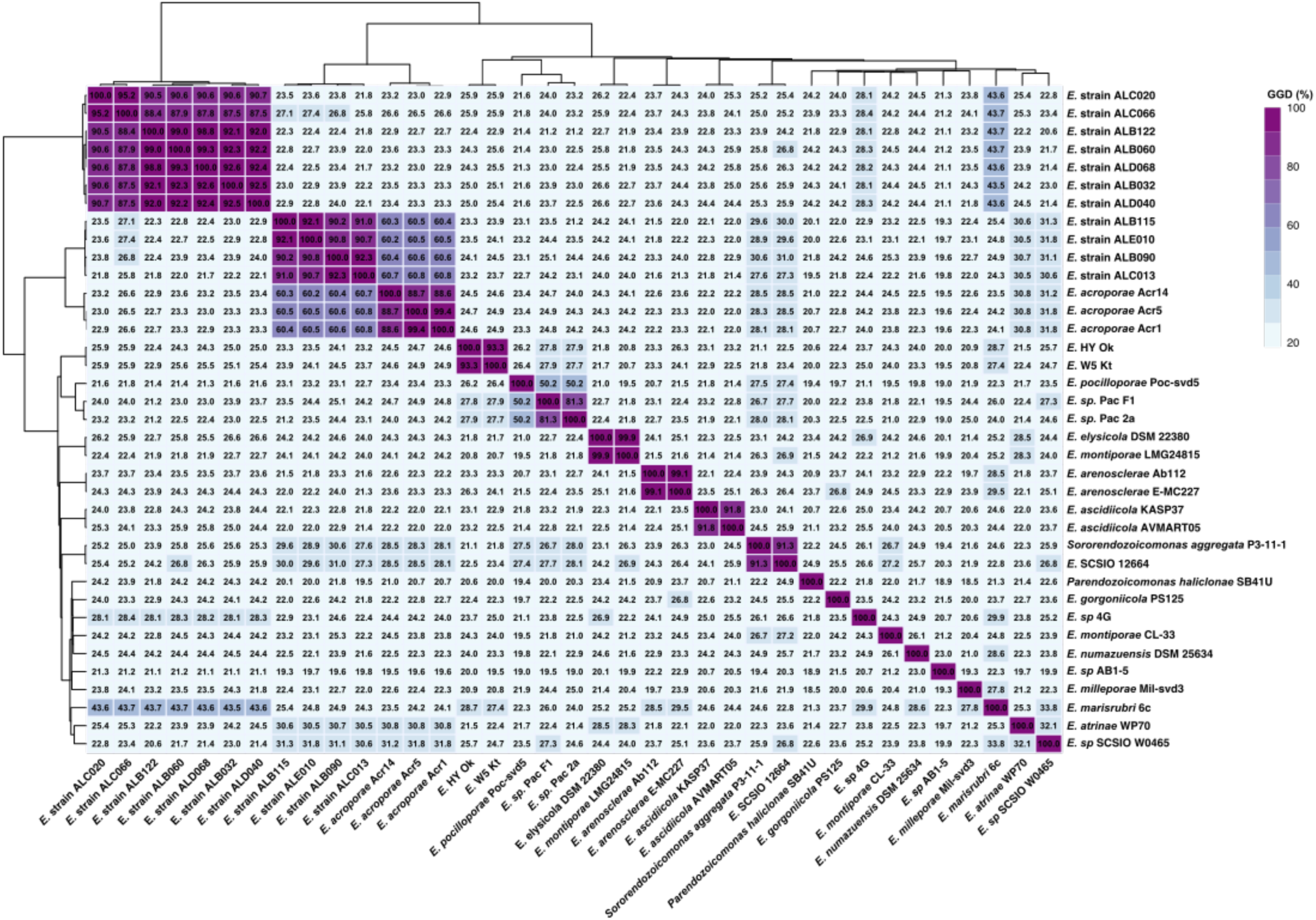
Heatmap showing the *in silico* GGD calculations between isolates obtained in this study and others from the literature.

**Figure S4.**
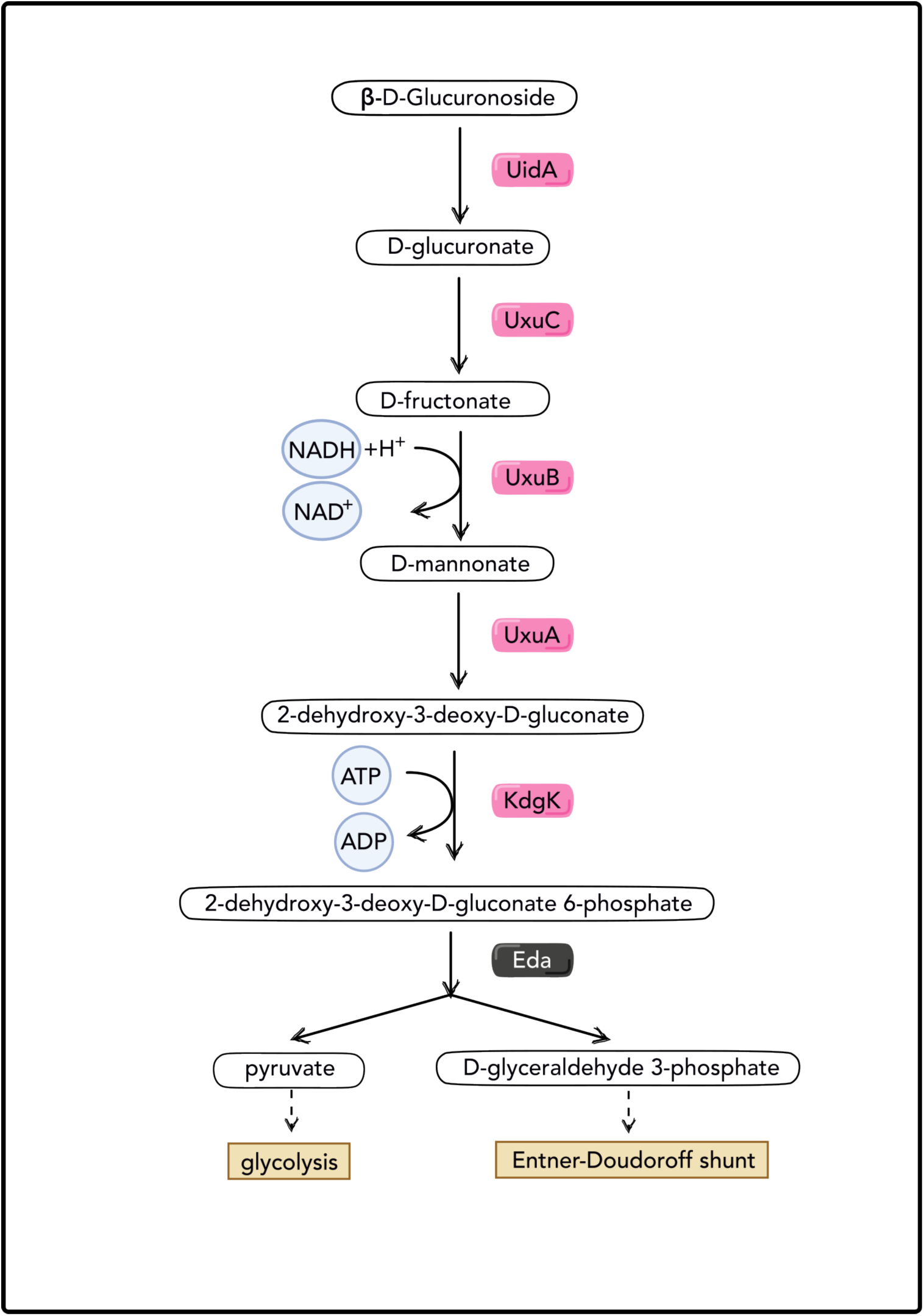
Glucuronate Interconversion. D-Galacturonate degradation pathway in bacteria responsible for the utilisation of glucuronate as a carbon source for energy, via UidA, UxuC, UxuB, and UxuA, KdgK. Proteins shown in magenta are exclusively found in Clade-A. Proteins shown in black are shared between Clade-A and Clade-B.

**Figure S5:**
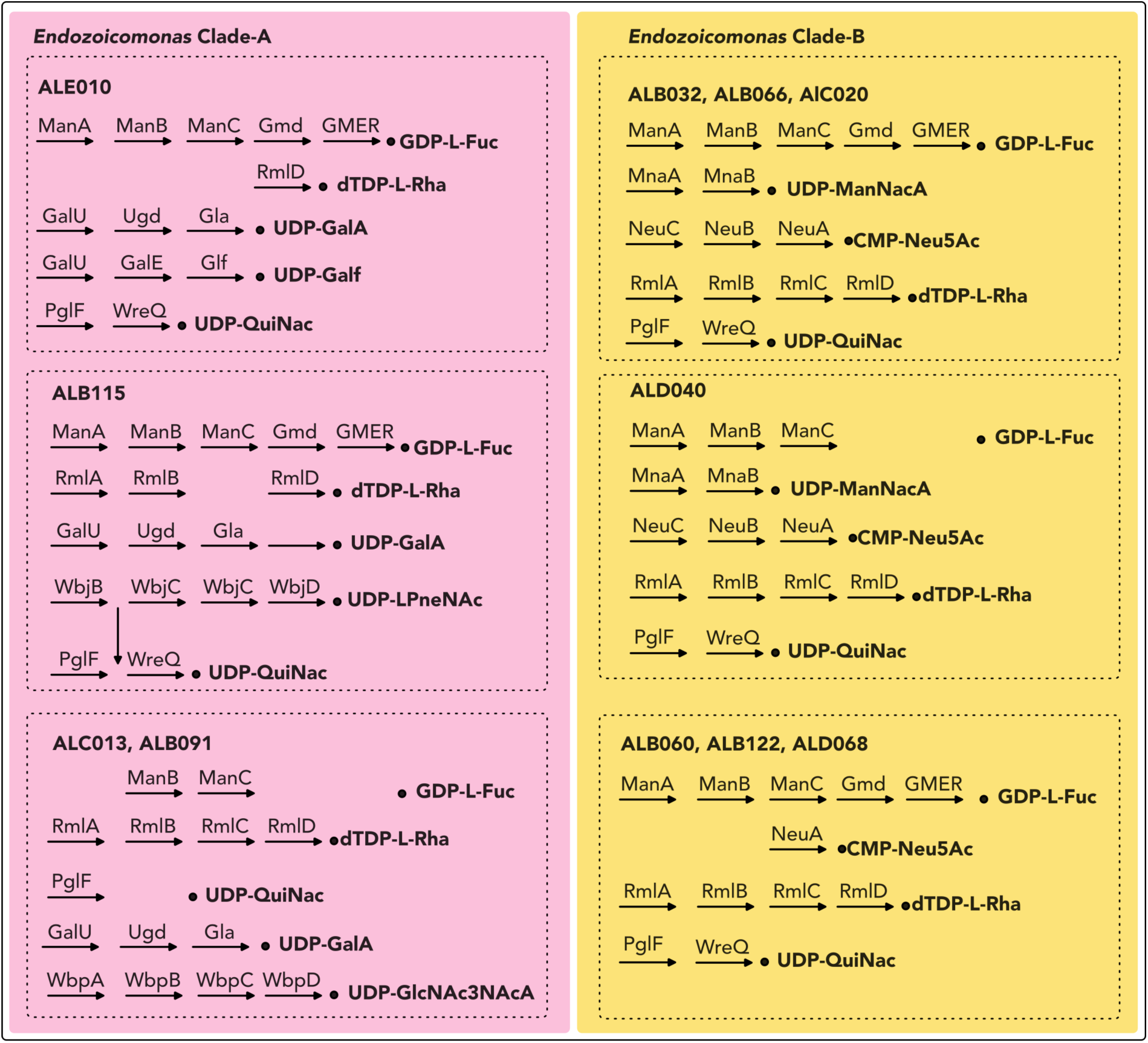
Comparison of O-antigen biosynthesis pathways in *Endozoicomonas* Clade-A and Clade-B. Clade-A strains are shown on the left, while Clade-B strains are depicted on the right. Pathways depicted in black represent fully complete pathways, while those with missing components indicate partial, incomplete, or absent pathways. Strain-specific differences between the two clades include the potential biosynthesis of UDP-N-acetylmannosamine (UDP-ManNAc), GDP-L-fucose (GDP-L-Fuc), and CMP-N-acetylneuraminate (CMP-Neu5Ac/sialic acid) in Clade-B. In contrast, all Clade-A strains possess the pathway for UDP-α-D-galacturonic acid (UDP-GalA), where galacturonic acid is a uronic acid derivative of galactose. Additionally, strains ALC013 and ALB091 contain the LPS gene cluster homologous to the Wbp pathway in *Pseudomonas aeruginosa*, which encodes enzymes required for the de novo biosynthesis of UDP-2,3-diacetamido-2,3-dideoxy-alpha-D-glucuronic acid (UDP-GlcNAc3NAcA), typically found in the O-antigen portion of the lipopolysaccharide layer in the Gram-negative cell wall.

**Figure S6:**
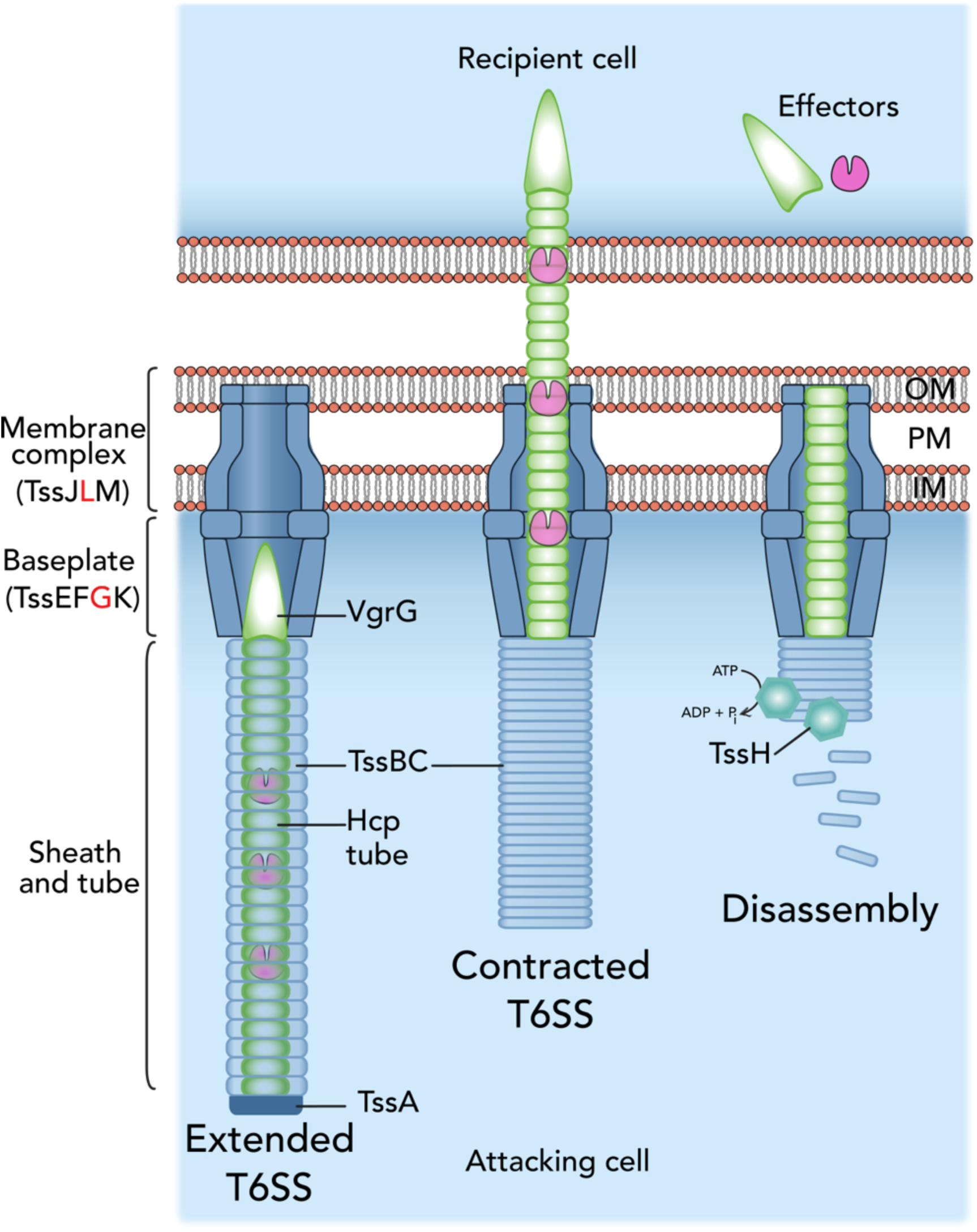
Representation of the T6SS gene cluster present in Clade-A The figure illustrates the roles of core T6SS components found in Clade-A, from assembly to effector delivery. The T6SS assembles with the baseplate (TssEFGK) and VgrG-PAAR spike docking onto the membrane complex (TssJLM), followed by contraction of the TssBC sheath, which propels the Hcp-VgrG complex into target cells. Upon contraction, the Hcp–VgrG neddle-like structure is driven into the target cell, releasing effectors. Subsequent disassembly involves the ATPase TssH, allowing the system to reset or relocate for another round of delivery. Proteins present in all Clade-A genomes are shown in black, while missing proteins are highlighted in red. OM: Outer membrane; PM: Periplasm; IM: Inner membrane.

**Figure S7:**
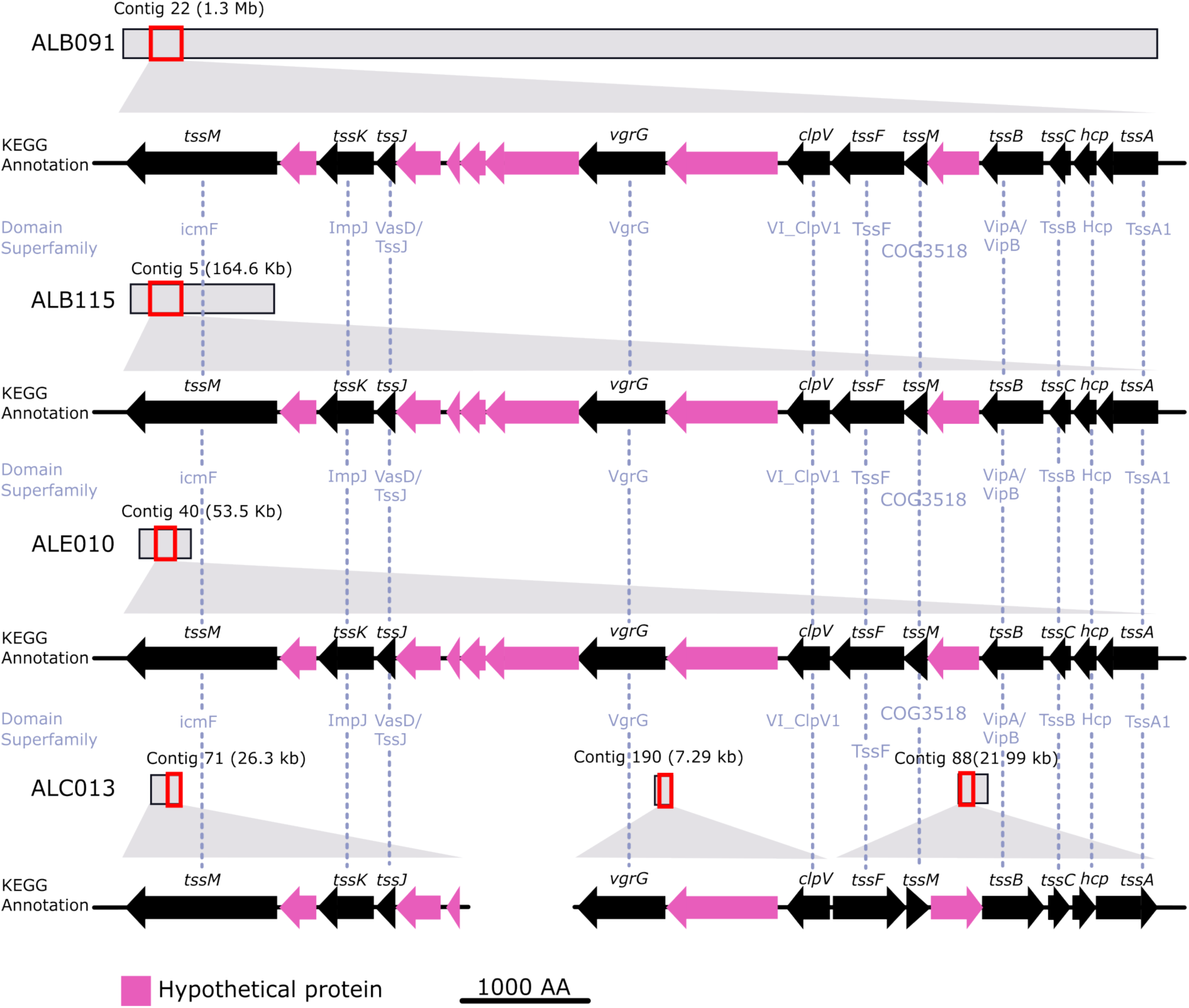
Illustration of the type VI secretion (T6SS) gene clusters present in Clade-A. Gene clusters are represented with the functional annotations depicted in different colors: black (proteins annotated with KEGG), magenta (hypothetical proteins based on RAST), and blue dotted line (annotated proteins through domain search).

**Figure S3-8:**
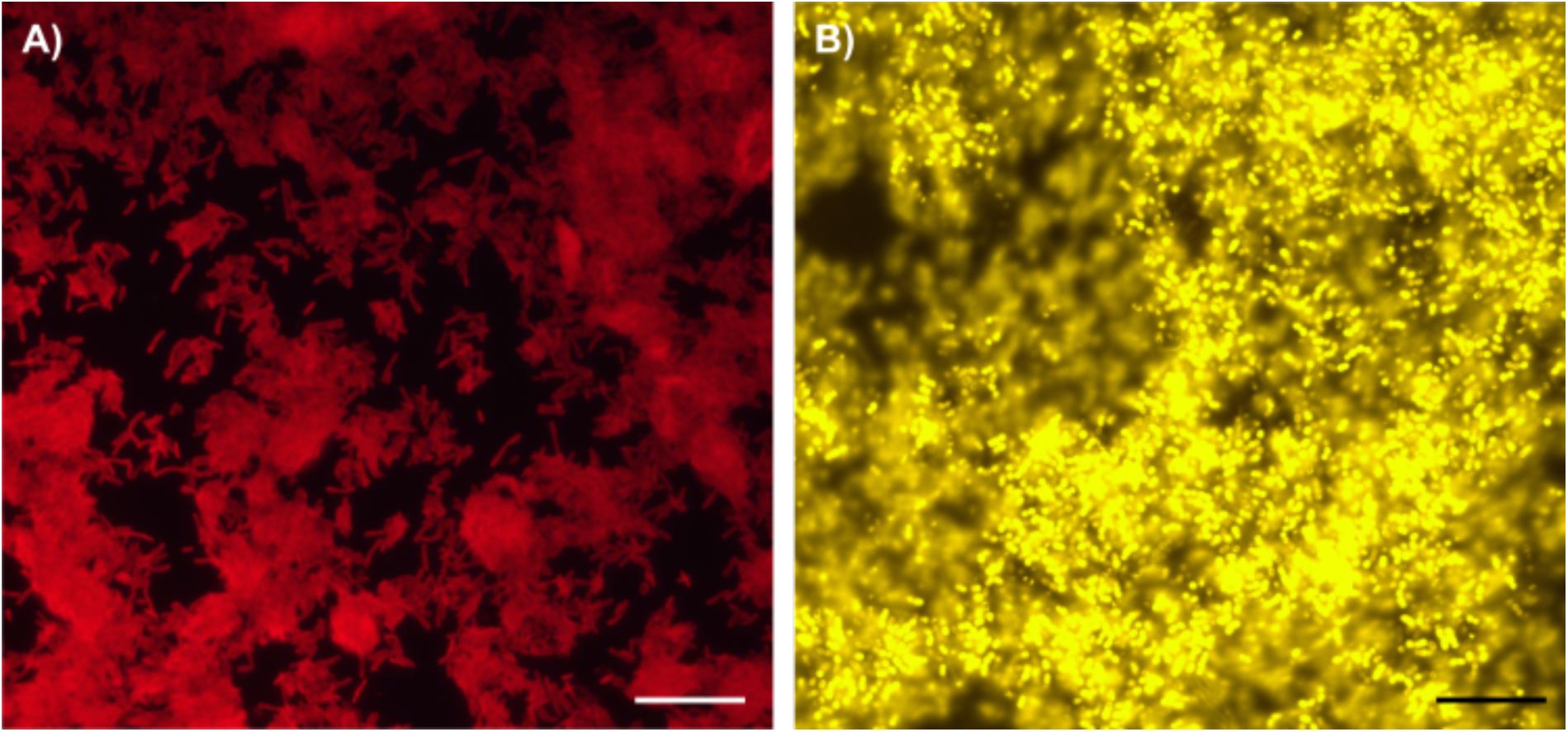
Clade-specific morphological difference in culture. In (A), bacterial isolates grown in liquid culture are stained with a Clade-A specific probe (Atto647), while in (B), isolates are stained with a Clade-B specific probe (Atto550). Clade-A cells (A) appear as single, rod-shaped bacteria, whereas Clade-B (B) secretes extracellular polysaccharide substances, possibly indicating differences in capsule formation. Scale bars represent 20 μm.

